# Profiling trace organic chemical biotransformation genes, enzymes and associated bacteria in microbial model communities

**DOI:** 10.1101/2024.03.25.586518

**Authors:** Lijia Cao, Sarahi L. Garcia, Christian Wurzbacher

## Abstract

Microbial biotransformation of trace organic chemicals (TOrCs) is an essential process in wastewater treatment for eliminating environmental pollution. Understanding of TOrC biotransformation mechanisms, especially at their original concentrations, is important to optimize treatment performance, whereas our current knowledge is limited. Here we investigated the biotransformation of seven TOrCs by 24 model communities. The genome-centric analyses unraveled the biotransformation drivers concerning functional genes and enzymes and responsible bacteria. We obtained efficient model communities for complete removal on ibuprofen, caffeine and atenolol, and the transformation efficiencies for sulfamethoxazole, carbamazepine, trimethoprim and gabapentin were 0-45%. Biotransformation performance was not fully reflected by the presence of known biotransformation genes and enzymes. However, functional similar homologs to existing biotransformation genes and enzymes (e.g., long-chain-fatty-acid-CoA ligase encoded by *fadD* and *fadD13* gene, acyl-CoA dehydrogenase encoded by *fadE12* gene) could play critical roles in TOrC metabolism. Finally, we identified previously undescribed degrading strains, e.g., *Rhodococcus qingshengii* for caffeine, carbamazepine, sulfamethoxazole and ibuprofen biotransformation, and potential transformation enzymes, e.g., SDR family oxidoreductase targeting sulfamethoxazole and putative hypothetical proteins for caffeine, atenolol and gabapentin biotransformation.

## Introduction

Trace organic chemicals (TOrCs) (e.g., pharmaceuticals, pesticides, and personal care products) discharged from industries, agriculture, hospitals and households have been frequently detected in natural water sources^1,2^. The increasing occurrence and accumulation of these ecologically harmful compounds has promoted worldwide research on TOrC removal technologies^3,4,5^. In general, biological treatment is an effective, economic and energy-saving strategy compared with conventional activated sludge and advanced oxidation processes^6,7^. Many studies have investigated the microbially mediated degradation of TOrCs and optimized its application in the water and wastewater treatment. For instance, Müller et al.^8^ introduced a novel approach of sequential biofiltration for the advanced treatment of secondary effluent and the pilot-scale experiments confirmed an increased removal of several TOrCs. Edefell et al.^9^ designed a novel process to increase biofilm growth in tertiary moving bed biofilm reactors by providing additional substrate from primary treated wastewater, which significantly improved TOrC removal.

Microorganisms play a vital role in the elimination of TOrCs via sorption and biodegradation (full mineralization) or biotransformation (incomplete removal of the parent compound)^10^. Biotransformation is considered as the primary removal process for most TOrCs in wastewater treatment plants, which is mainly attributed to oxidative reactions and ammonia oxidizing bacteria^11,12^. There have been many studies focusing on pure cultures or synthetic microbial consortium that attempted to decipher the metabolic mechanisms including biotransformation byproducts and pathways, key metabolic enzymes, and interspecies interactions^13,14,15^. Nevertheless, our current knowledge about biotransformation at the level of gene/enzyme-chemical interactions is still limited especially for refractory TOrCs. Addressing this issue is critical for the targeted optimization of biological treatment processes which requires a better understanding of the microbial functionality.

In our previous study on the establishment of TOrC-degrading model communities via environmental subset, we obtained eleven taxonomically non-redundant cultivated model communities with different removal abilities on TOrCs^16^. Since model communities are well-defined and standardized, they are great tools to identify key driving agents of TOrC biotransformation, i.e., responsible genes and enzymes, and associated microorganisms as well as their functions in the community. However, we observed in our previous study that living in the environment consisting of 27 TOrCs, the biotransformation of individual chemical by model communities was not high-effective and the majority of TOrCs remained unremoved. A hypothesis is that various TOrCs with different physicochemical properties complicate their efficient biotransformation^17^. Accordingly, to achieve TOrC-specific promising degraders, here we selected seven TOrCs that are frequently detected in natural water sources, i.e., atenolol, caffeine, carbamazepine, gabapentin, ibuprofen, trimethoprim and sulfamethoxazole. We then transferred 6 model communities from our previous study^16^ and cultivated *de novo* 18 model communities with the seven chemicals as the substrate either with single chemical or with different mixtures. We obtained the metagenomes of the 24 model communities and comparatively examined (i) the microbial removal performances on the seven TOrCs, (ii) the potential genes and enzymes responsible for the initial metabolic step, and (iii) associated bacteria and the roles they play in the community. The aim of this study is to correlate TOrC biotransformation efficiencies with the presence of currently reported responsible genes and enzymes, and to mine putative novel functions and degraders to expand our knowledge. We hypothesize that TOrC biotransformation efficiencies could be mirrored by the related functional genes and corresponding enzymes. The genome-centric analyses based on these simplified model communities reinforce our understanding of TOrC biotransformation by complementing existing knowledge. Our research could further benefit for designing new approaches for engineering microbes with enhanced biotransformation abilities such as assembly of pathways using enzymes from diverse bacteria to bioremediate TOrC-contaminated environments.

## Materials and methods

### TOrC selection

Seven TOrCs, atenolol, caffeine, carbamazepine, gabapentin, ibuprofen, trimethoprim and sulfamethoxazole, were selected in this study. These seven TOrCs represent frequently occurring pollutants in municipal wastewater derived with different degree of biodegradability (Table S1). Caffeine and ibuprofen are easy to be biotransformed, and their transformation mechanisms are relatively well documented^18,19^. Carbamazepine, gabapentin and trimethoprim are rather persistent and poorly biodegradable, and few studies have addressed their biotransformation^20,21,22^. Atenolol and sulfamethoxazole are reported to be fully or partially degraded by microbial strains^23,24^. The uses, biotransformation efficiencies, occurrence in aquatic systems, and ecological risks can be found in Table S1.

### Model community cultivation

Six model communities were obtained from our previous experiments^16^. They were enriched by a mixture of 27 TOrCs and showed biotransformation abilities on the seven investigated compounds. Moreover, we followed the model community establishment workflow that we described before and cultivated another 18 communities. In brief, five samples collected from tap water (Garching, Germany), technical sand treated with tertiary effluent (Garching, Germany), soil (Garching, Germany), surface and deep sediment (47°47′16“N, 11°18′16”E, Osterseen, Germany) on November 2021 were enriched by the seven TOrCs (either individually or jointly, 11 treatments) for six months, resulting in 165 treatment bottles (each TOrC treatment was conducted in triplicate) (Table S2) and 22 blank bottles (11 bottles without cells and 11 bottles with autoclaved cells). The blanks were set as controls to assess the abiotic removal of TOrCs. The enrichment process was divided into six phases with TOrC concentrations set as 50 nmol/L (P1), 250 nmol/L (P2), 500 nmol/L (P3), 1000 nmol/L (P4), 2500 nmol/L (P5), 50 nmol/L (P6), respectively. Cell counts were determined by the flow cytometer at the end of each phase to ensure the cell growth following^16^. After the enrichment, microbial communities from P6 were then subjected to the dilution-to-extinction step for reducing the species richness. Based on our previous results that enriched microbial communities achieved successful growth above 1×10^6^ cells/mL (the ubiquitous density observed in aquatic environment) at the dilution threshold of 10 cells/mL, here we diluted the samples from P6 (pooled triplicates) to 10 cells per well in 96 deep well plates (PP, SARSTEDT, Germany) which contained 1 mL growth medium per well. The growth medium consisted of 5 nmol/L TOrCs and mineral salts as described in our previous study^16^. Diluted microbes were incubated in the plates in the dark at room temperature for three weeks followed by the measurement of growth. Then, 500 μL suspension of successful growth wells were transferred to new plates containing 1.5 mL medium per well and were cultivated for two weeks, to get enough volume for the subsequent TOrC removal determination and DNA extraction. Taxonomy refinement was achieved based on the 16S rRNA sequencing (same procedure following Cao et al.^16^) and taxonomic assignment by Emu^25^. The treatment conditions and the preliminary selection of model communities were listed in Table S2.

To evaluate the removal rate of each TOrC (not only the feeding TOrC since cultivation) by the 24 model communities, 100 μL suspension of the obtained 24 model communities were added into 1 mL medium spiked with 5 nmol/L individual TOrC, and were incubated for three weeks.

### Determination of TOrC biotransformation

TOrC reduction was determined using liquid chromatography coupled with tandem mass spectrometry (LC–MS/MS). An ultra-high performance liquid chromatograph (PLATINblue UPLC, Knauer, Germany) was equipped with a Phenomenex Kinetex PFP 100-Å chromatographic column (150 × 3 mm, 2.6 μm). The UPLC was connected to a Turbo V ion source of a triple quadrupole mass spectrometer (Triple Quad 6500, SCIEX, USA) operated in positive and negative electrospray ionization mode. A binary gradient system was applied, consisting of mobile phase A, Milli-Q water with 0.1% formic acid, and mobile phase B, LC-MS grade acetonitrile (Merck, Germany). Prior to the measurement, 1 mL of samples collected from model community cultivation medium were diluted 2 times to obtain enough volume. 1900 μL of diluted samples were spiked with 100 μL of internal standard, and then filtered through 0.22 μm polyvinylidene difluoride (PVDF) membrane filters into 2 mL amber glass vials for injection. The internal standard method was used for quantification. Isotope-labeled internal standards were available for all analytes. Standard samples at concentrations of 2.5, 5, 10, 25, 50, 100, 250, 500, 1000, 2500, 5000, 10000 ppt were prepared for calibration curves. The data processing was performed on MultiQuant software.

### Metagenome sequencing, assembly, binning and taxonomy classification of MAGs

DNA was extracted from 24 model communities (600 μL of each community) using the DNeasy PowerSoil Pro Kit (QIAGEN) according to manufacturers’ instruction. Extracted DNA was quantified using dsDNA Broad Range Assay (DeNovix, USA) in a fluorometer (DeNovix, USA). The metagenomic sequencing was performed on PromethION P24 (Oxford Nanopore Technologies) with the R10.4.1 flow cell at LMU Gene Center (Munich, Germany). 100 ng genomic DNA of sample in 12.5 μL was subjected to prepare a total of 75 μL library pool using native barcode ligation kit 96 (v14). The average sequencing depth was about 3 Gb per model community. Obtained raw reads were firstly demultiplexed according to their barcodes using Guppy v3.6.0 (https://timkahlke.github.io/LongRead_tutorials/BS_G.html). Demultiplexed fast5 reads were duplex-basecalled and converted to FASTQ files using Dorado v0.3.2 (https://github.com/nanoporetech/dorado/). Barcodes and adapters were also trimmed by Dorado. All sequenced reads from one sample were merged followed by quality control using Filtlong v0.2.1 (https://github.com/rrwick/Filtlong) with the parameters “--min_length 1000 --min_mean_q 10”. Filtered clean reads of each metagenome were assembled and binned by NanoPhase v0.2.3^26^. Specifically, metaFlye v.2.9-b1768^27^ was used to assemble trimmed long reads with the option “--nano-hq -i 5 -g 4m”. Afterwards, draft metagenome assembled genomes (MAGs) were constructed using MetaBAT2^28^ and MaxBin2^29^, and were subjected to the refinement step conducted by MetaWRAP v1.3.2^30^ to retain the best representative and non-redundant MAGs (completeness > 50% and contamination < 10%). Finally, long reads were mapped to the draft MAGs using minimap2 v2.21-r1071^31^ with 90% identity and coverage to produce clusters. Draft MAGs were then polished based on the clusters with one round of Racon v1.4.22 (https://github.com/isovic/racon) and one round of medaka v1.4.3 (https://github.com/nanoporetech/medaka) to generate high-quality final MAGs. The relative abundance of each MAG was evaluated by SingleM v0.16.0^32^. The taxonomies of all MAGs derived from 24 model communities were classified by GTDB-Tk v2.3.2^33^. The phylogenetic tree of all MAGs based on 120 single-copy marker proteins for bacteria was constructed using the maximum likelihood method via FastTree v2.1.11^34^ and visualized by iTOL v5^35^.

### Identification of TOrC biotransformation genes, enzymes and pathways

Based on the published literatures, enviPath (a database and prediction system for the microbial biotransformation of organic environmental contaminants)^36^ and MetaCyc (a curated database of experimentally elucidated metabolic pathways from all domains of life)^37^, we collected information on the biotransformation pathways, related genes and enzymes, and degrading strains of the seven TOrCs, as shown in Figure 1 and Table S3. The functional annotation of MAGs was conducted by Prokka v1.14.5^38^ based on the amino acid sequences predicted by Prodigal v2.6.3^39^. For the genes and enzymes that were not documented in the Prokka annotation databases (e.g. UniProtKB, RefSeq and Pfam), we downloaded their amino acid sequences and identified their homologous proteins using orthoFind with the BLAST e-value as 5e-05, the minimal identity percent from BLAST alignment as 53%, and the minimum length for a domain region as 28^40^. The amino acid sequences were used in a BLAST search to find putative homologous protein. Then, a PSI-BLAST search was performed based on the query sequences and candidate protein sequences to find new and more complete homologous sequences in the Swiss-Prot database. All sequence alignments were produced using ClustalW, and the maximum likelihood trees with 1000 bootsrap were constructed using IQ-TREE v2.2.5^41^. The presence of biotransformation genes and enzymes and their homologs were searched against the annotation results of Prokka. Besides the known functional genes and enzymes, we conducted the comparative genomic analysis using OrthoFinder v2.5.5^42^ to identify orthologous groups across 24 model communities and correlate the orthogroups with TOrC removal mining for potential novel biotransformation enzymes. Amino acid sequences of MAGs were merged for each corresponding model community and the concatenated 24 genomes were used as input files in OrthoFinder. DIAMOND and MAFFT were used to the all-versus-all sequence search and multiple sequence alignment, respectively. Gene ontology (GO) functional annotation of the concatenated 24 genomes was performed by PANNZER^43^. Enrichment of GO terms present in each model community’s genes relative to the customized background consisting of all communities’ genes was performed using clusterProfiler^44^ in R with the function “enricher” and “compareCluster”, and significant enrichment was determined at an adjusted pvalue of 0.05.

**Figure 1.**
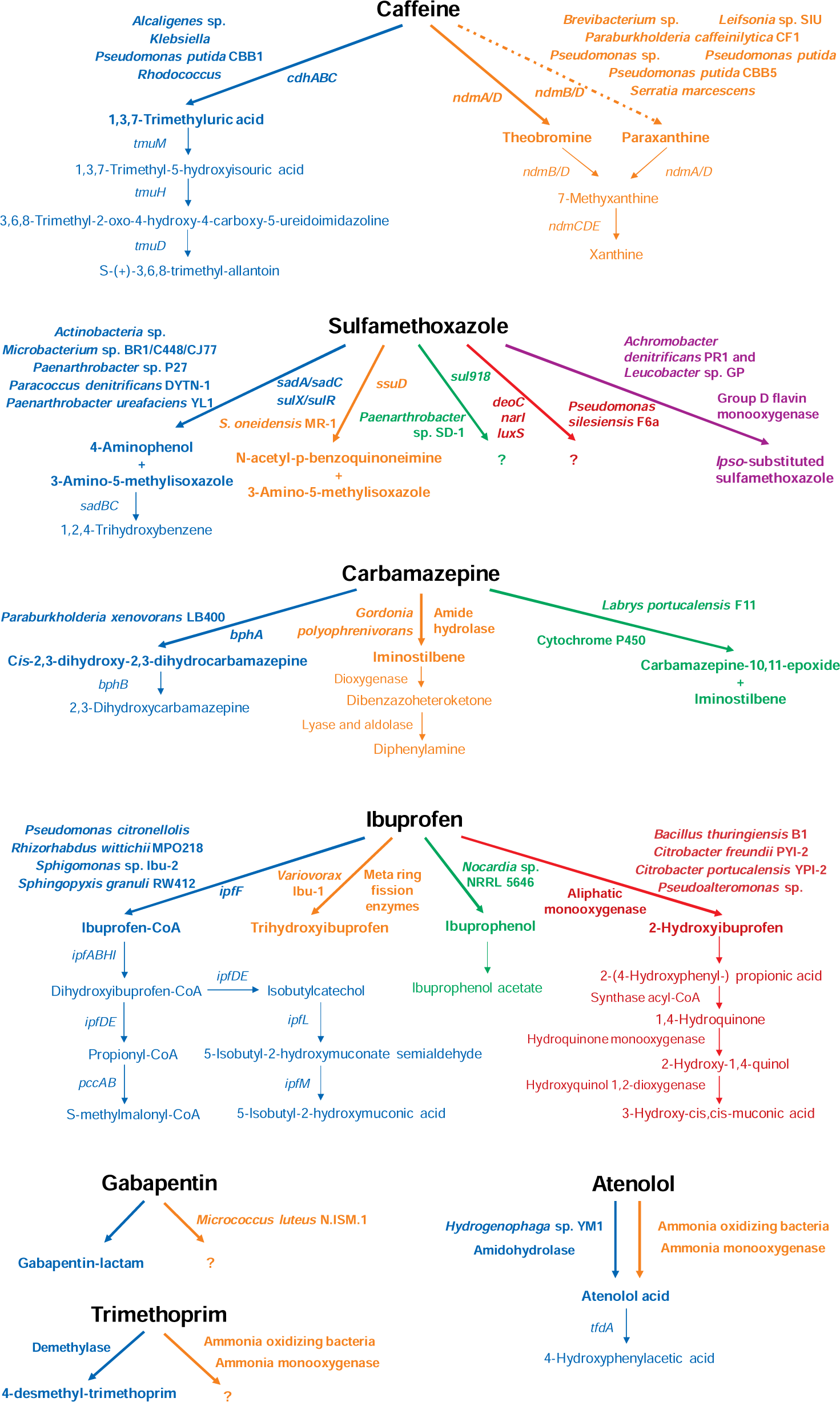
Overview on currently known biotransformation genes, enzymes, pathways, and associated bacteria of caffeine, sulfamethoxazole, carbamazepine, ibuprofen, gabapentin, trimethoprim, and atenolol. Only experimentally validated biotransformation information was included. Colors indicate different metabolic pathways. Bold arrows and text represent the first biotransformation steps and involved genes, enzymes and degraders which were mainly discussed in this study. Question marks represent unknown transformation products. Dash line in caffeine indicates the minor biotransformation pathway.

## Results and discussion

### TOrC biotransformation efficiencies

The 24 model communities exhibited different removal abilities on the seven TOrCs as shown in Figure 2. In general, most of the communities (19/24) can fully eliminate ibuprofen. While chemicals with aromatic rings are usually resistant to degradation by microorganisms, many studies indicate that ibuprofen is biodegradable despite of its structural characteristics^45^. In a WWTP aeration tank, greater than 95% of ibuprofen was removed with aerobic biodegradation being the dominant mechanism^46^. In a lab-scale cultivated consortium from sewage sludge, 100% of ibuprofen (1 mg/L) was degraded in solution in 6 h and 90% of ibuprofen (10 mg/kg) was degraded in sewage sludge in 16 days^47^. Sulfamethoxazole, carbamazepine and gabapentin can only be partially biotransformed by several model communities with the maximum percentage of 45%, 40% and 42%, respectively. These compounds are often refractive to biodegradation and the reported biological removal efficiencies in water and wastewater treatment are in general below 10%^20,48,49^. Their chemical structures (e.g., benzene and isoxazole ring in sulfamethoxazole, cyclohexane ring in gabapentin) and potential toxicity to microorganisms^50^ could retard the biotransformation. Caffeine has been reported to be an easily degradable compound with high removal efficiency in WWTP^51^, bioreactors^52^, managed aquifer recharge^23^ and diverse microbial strains^53^. Surprisingly, in the present study, only one model community E5 can degrade 100% of caffeine, and other degrading community (A2, A10 and G10) showed merely 22-27% removal, which may point to a preference of ibuprofen over caffeine. Atenolol was reported as a moderately biodegradable compound^23^ and trimethoprim is quite recalcitrant^54,55^. It is consistent in our study that there were three communities (C7, C9 and D8) degrading 85-100% of atenolol and the remaining communities showed none or slight (<27%) removal, while for trimethoprim the average removal efficiency was only 2% among the 24 model communities. Previous studies reported that the kinetics and efficiencies of TOrC biotransformation were impacted by TOrC initial concentration^56,57,58,59^. For example, pesticide in the low μg/L concentration range often lead to reduced biodegradation^60,61^. This effect was suggested to be substrate-specific^62^, which might explain the same concentration of the seven TOrCs resulted in different removal limits. Overall, ibuprofen, caffeine and atenolol can be 100% removed by high-efficient degrading model communities with ibuprofen being the most widely removed TOrC. Two model communities removed 23% and 40% of carbamazepine, three communities remove 22-42% of gabapentin, and five communities removed 21-45% of sulfamethoxazole. Trimethoprim was the most persistent chemical that no model community could remove. The community E5 was the best degrader with the ability of biotransforming five TOrCs (i.e., caffeine, ibuprofen, sulfamethoxazole, carbamazepine and gabapentin).

**Figure 2.**
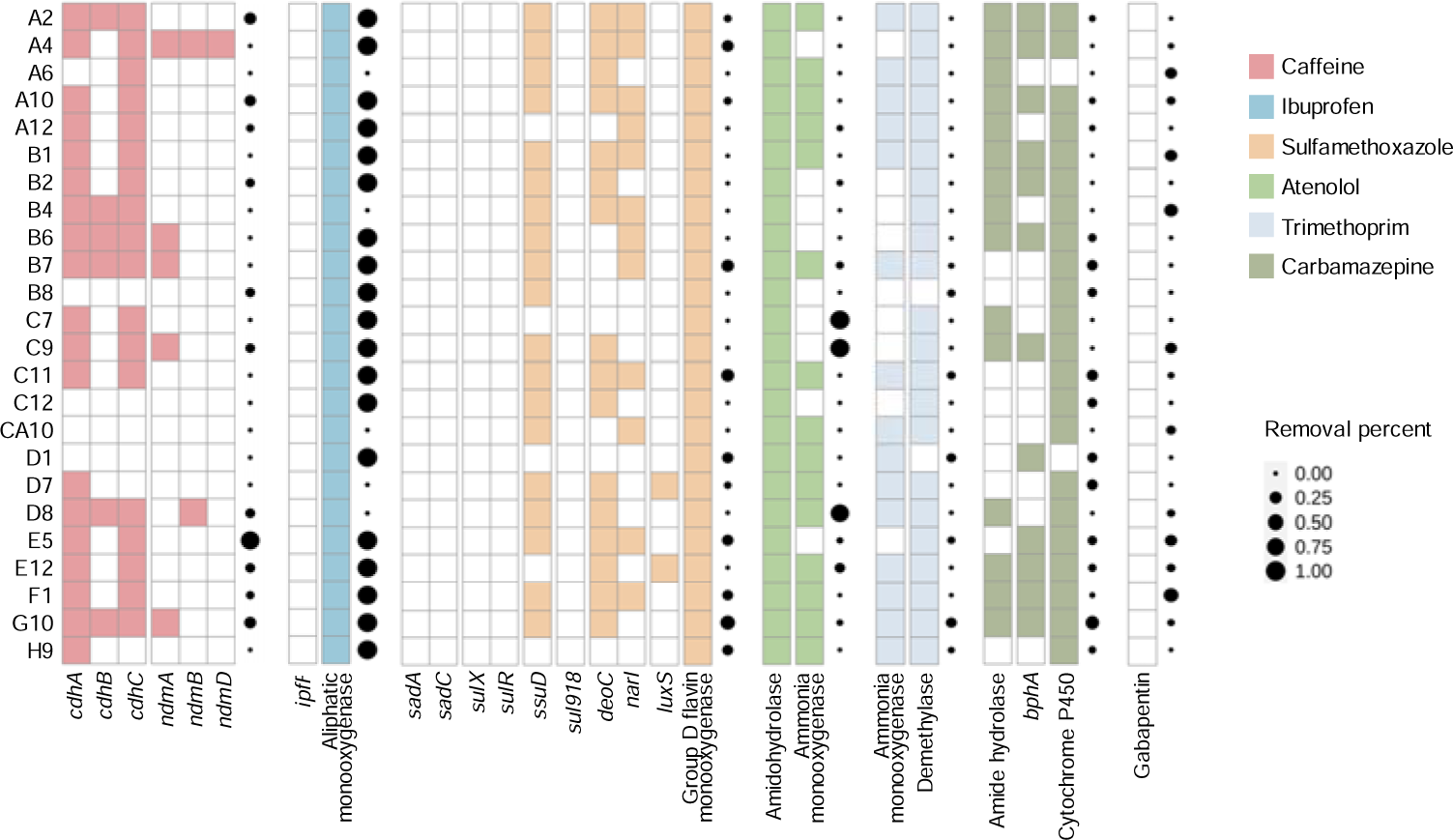
TOrC biotransformation efficiencies and the presence/absence of first-step biotransformation genes and enzymes in the 24 model communities. Colors represent presence, blank represents absence. The dot plot indicates the removal percentage. No biotransformation genes or enzymes for gabapentin have been reported yet.

### Presence of biotransformation genes and enzymes in model communities

To indicate TOrC removal performance by functional genes and enzymes, we identified the presence/absence of biotransformation genes in the metagenomic annotation results of model communities based on the previous literatures, enviPath and MetaCyc (Figure 2, Table S3). These collected genes and enzymes included first-step biotransformation genes involved in caffeine, ibuprofen, sulfamethoxazole and carbamazepine, and first-step biotransformation enzymes involved in ibuprofen, sulfamethoxazole, atenolol, trimethoprim, carbamazepine and gabapentin. We hypothesized that TOrC biotransformation efficiencies could be mirrored by the related functional genes and enzymes. There were 20 model communities possessing at least one of the *cdhABC* genes responsible for caffeine dehydrogenation (Figure 2). The *ndmA*, *ndmB* and *ndmD* genes involved in caffeine demethylation were present in six model communities (i.e., A4, B6, B7, C9, D8 and G10). Community B8, C12, CA10 and D1 which did not have any caffeine biotransformation genes showed no removal of caffeine. However, not all communities containing related genes could biotransform caffeine, and the best performer E5 only contained *cdhA* and *cdhC* genes which also appeared in other non-degrading model communities (e.g., B1, C7 and C11). It also happened in ibuprofen, sulfamethoxazole, atenolol, trimethoprim and carbamazepine that not all communities having the biotransformation genes and enzymes showed removal on corresponding chemicals, and the same distribution of genes did not indicate similar removal efficiency. For example, aliphatic monooxygenase that metabolize ibuprofen to 2-hydroxyibuprofen was present in all communities, while A6, B4, CA10, D7 and D8 did not reduce any ibuprofen. For gabapentin, although there have been no biotransformation genes or enzymes reported yet, the removal in our study indicated the possibility of previously undescribed functions. The *ipfF*, *sadA*, *sadC*, *sulX*, *sulR* and *sul918* genes were not annotated by Prokka so that they were absent in all communities. In the following section, we identified their homologs to find the potential correlations with TOrC removal.

In summary, our hypothesis was partially rejected since the presence of biotransformation genes and enzymes did not fully reflect TOrC removal efficiencies by model communities. The threshold concentration effect (that is, the lowest substrate concentration below which no appreciable growth of specific degrader organisms could be observed that leads to no reduction of substrate)^63,56^, was suggested to result in the lack of induction of biotransformation gene expression and enzymatic activities^64,65^. This could also explain the inconsistence between gene presence and not finding corresponding TOrC removal in our study, since the metagenomic analysis shows gene presence but that does not guarantee gene expression. Further metatranscriptomic analysis of functional gene expression patterns is suggested to obtain insights into deciphering the relationships between TOrC presence and the regulation of biotransformation genes and eventually removal performance.

### Phylogenetic analysis of biotransformation genes and enzymes and their homologs

The *ipfF*, *sadA*, *sadC*, *sulX, sulR* and *sul918* genes (responsible for ibuprofen and sulfamethoxazole biotransformation) were not annotated by Prokka, however, it is likely that these genes may have diversified in microbial lineages by vertical evolution. Thus, the five genes above were used as queries to identify potential homologs using orthoFind. Biotransformation genes for caffeine are well documented so that they were not investigated here. For atenolol, gabapentin and trimethoprim, their biotransformation genes have not been discovered and are not able to serve as reference sequences. The number of homologs, functions, domain architecture of the query sequences were listed in Table S4. Maximum likelihood trees were constructed for the IpfF, SadA, SadC and Sul918 proteins to further investigate the evolutionary links among the various homologs (Figure S1).

There were 59 homologs to IpfF (ibuprofen CoA ligase) among which CaiC (crotonobetaine/carnitine-CoA ligase), AscA (acetyl-coenzyme A synthetase), MenE (2-succinylbenzoate-CoA ligase), YhfT (uncharacterized acyl-CoA ligase), FadK (medium-chain-fatty-acid-CoA ligase), LcfB (long-chain-fatty-acid-CoA ligase), FadD3 (long-chain-fatty-acid-CoA ligase), FadD13 (long-chain-fatty-acid-CoA ligase), FadD (long-chain-fatty-acid-CoA ligase) and RpfB (fatty Acyl-CoA ligase) involved in the ligase activity were present in the 24 model communities (Figure S1a, Figure 3a). The long-chain-fatty-acid-CoA ligase is known to converting xenobiotics by targeting the carboxyl or hydroxyl groups as the initial metabolic step. For instance, Harb et al.^66^ found this enzyme facilitated the biodegradation of two hydroxyl-containing micropollutants (atenolol and acetaminophen) in an anaerobic MBR system. Pirete et al.^67^ identified the long-chain-fatty-acid-CoA ligase as the main enzyme involved in diclofenac biotransformation.

**Figure 3.**
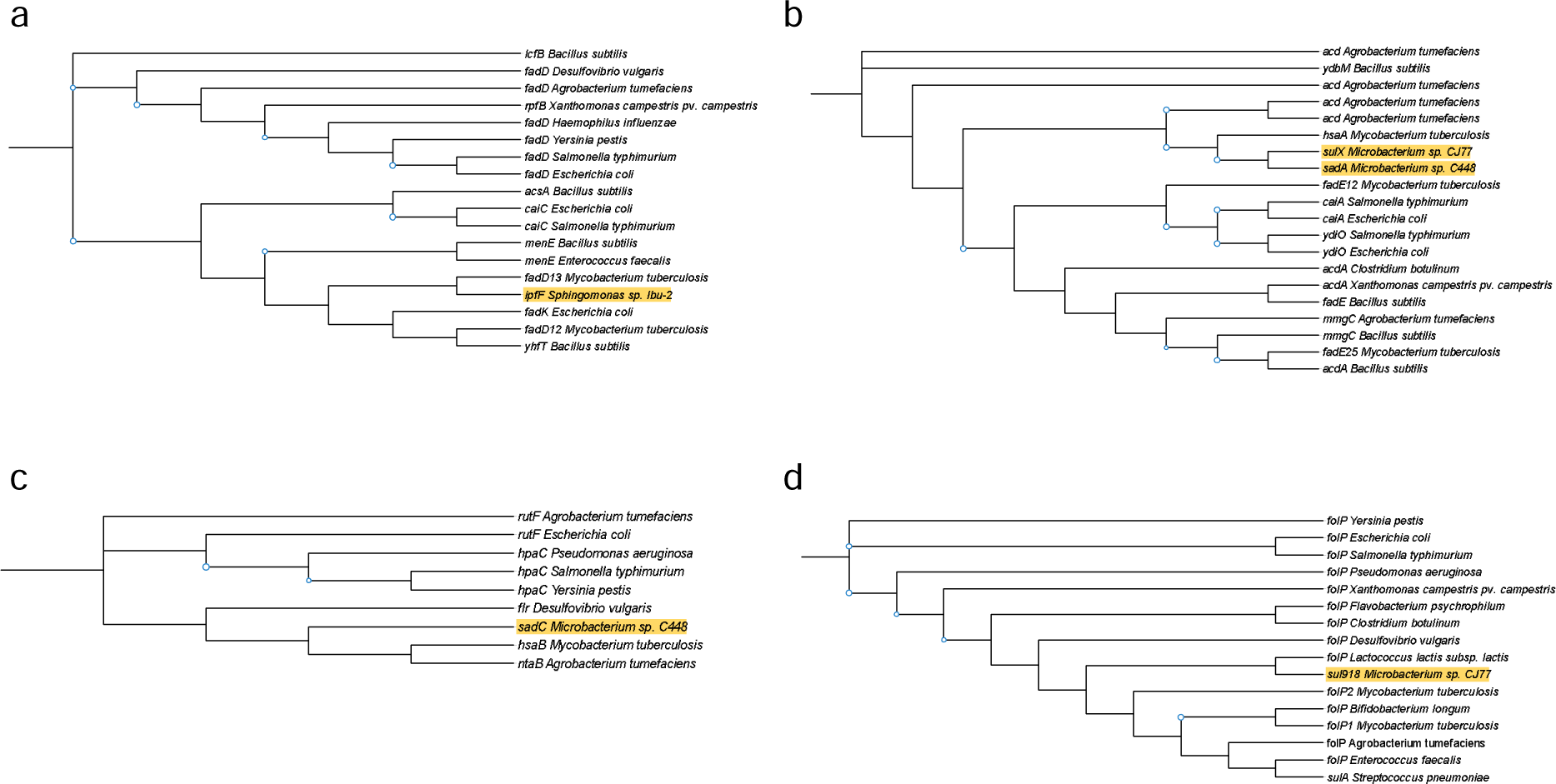
Phylogenetic trees of (a) ipfF, (b) sadA, (c) sadC, and (d) sul918 genes and their homologs present in model communities. Homologous proteins were identified based on the amino acid sequences of these biotransformation genes using OrthoFind. Phylogenetic trees were constructed using maximum likelihood method by IQ-TREE and modified by iTOL. Bootstrap support values above 85 are indicated at node.

There were 82 homologous proteins to SadA (sulfonamide monooxygenase) with the function of flavin adenine dinucleotide binding and acyl-CoA dehydrogenase activity which was the same as SulX (Figure S1b, Figure 3b). SulX has been proven to be a homologous protein to SadA^68^. The *hsaA* gene encoding a flavin-dependent monooxygenase located most closely to *sadA* gene among the homologs that can be identified in the model communities. The flavin reductase SadC had 47 homologs, HpaC (4-hydroxyphenylacetate 3-monooxygenase reductase component), RutF (FMN reductase), NtaB (FMN reductase), HsaB (flavin-dependent monooxygenase, reductase subunit) and Flr (flavoredoxin) were present in the model communities (Figure S1c, Figure 3c). It was the same to SulR having the identical amino acid sequences with SadC. SadA and SadC were responsible for the initial cleavage of sulfonamides^24^, and the gene cluster *sulX* and *sulR* containing homologs of SadA and SadC^68^ was also reported to degrade sulfonamides. SadA is highly specific to catalyze the ipso-hydroxylation of sulfamethoxazole releasing 4-aminophenol, while the auxiliary role of SadC in electron transport can easily be replaced by other enzymes with similar function^24^, indicating the predominant role of SadA in the initial step of sulfamethoxazole biotransformation.

The sulfonamide resistant gene *sul918* had most of the homologs characterized as *folP* encoding dihydropteroate synthase (Figure S1d, Figure 3d). The sulfonamide resistance mechanisms are mediated by mutations in *folP* and/or acquisition of *sul* genes. To date the origins of *sul* genes are not clear, but a recent study suggested that the *sul* genes evolved from lateral transfer of chromosomal *folP* genes^69^. Although *sul918* is not the sulfamethoxazole degrading gene, its co-occurrence with *sad* gene cluster was found to favor the efficient degradation of sulfamethoxazole^70^. The possibility of antibiotic resistance genes (e.g., *sul1*, *sul918*) conferring antibiotic biotransformation might indicate the mobilization of biotransformation genes associated with mobile genetic elements between different taxa. Contradictory finding was also reported that a *Paenarthrobacter* strain containing a complete *sad* gene cluster and *sul* genes (*sul918* and *sul1*) displayed limited sulfonamide removal^71^. Additional studies are required to decipher the relationships between sulfonamide resistance and biotransformation, which is critical for understanding the dissemination of antimicrobial resistance.

The known biotransformation genes and enzymes together with their homologs provided us with more comprehensive understanding of the function of members of the same protein family for TOrC biotransformation. Phylogenetic analysis of homologs also facilitates the discovery of potential novel biotransformation genes^72^. In many cases, the presence of functional similar homologous proteins acted as surrogates to initiate the metabolic reactions^68,73,74^. This further points to the possibility of functional gene diversity as an important driver for the transformation of some compounds^75^. However, the homologous proteins often have differences in structure, such as the substrate binding pocket, which could influence TOrC binding to the active site of the enzyme and therefore influence the biotransformation efficiency. For example, SadA had a wider pocket than its homolog XiaF, and *Leucobacter* strain GP containing SadA showed better sulfonamide removal than *Microbacterium* sp. BR1 containing XiaF^76^.

### Potential biotransformation pathways and associated bacteria carrying related genes and enzymes

To characterize the specific MAGs carrying the biotransformation genes and enzymes, we recovered in total 88 high-quality draft genomes from the 24 model communities’ metagenomes (abundance, completeness and contamination are provided in Table S4). Each model community consisted of 1-8 MAGs. According to the Genome Taxonomy Database (GTDB), a total of six phyla were identified with the most abundant phyla being Proteobacteria (nLJ=LJ64) and Actinobacteriota (nLJ=LJ16) (Figure 4). Thirty-two of these MAGs were classified to the species level, four MAGs were classified to the family level, and the remaining MAGs were identified to the genus level, indicating novel taxa at different taxonomical levels. Using GTDB nomenclature that newly delineated and uncultured taxa are allocated with alphanumeric placeholder names, we found eighteen MAGs were assigned to genus or species with such placeholder labels (e.g., g_62-47, g_DSPA01, s_Afipia sp017474385, s_Variovorax sp900115375) (Figure 4). Such taxonomic novelty with 75% of MAGs affiliated with new classification is likely driven by niche adaptation to the distinctive cultivation environment where TOrCs served as the sole carbon source^77,78^. Accordingly, the utilization of model communities adapted to TOrCs offers us an opportunity to target key microorganisms that are easily submerged in natural microbiome and to uncover their functions that are barely addressed. The frequencies of biotransformation genes and enzymes including their homologs in each MAG of TOrC-degrading model communities were determined to identify the main microbial players in the initial biotransformation step (Figure 5).

**Figure 4.**
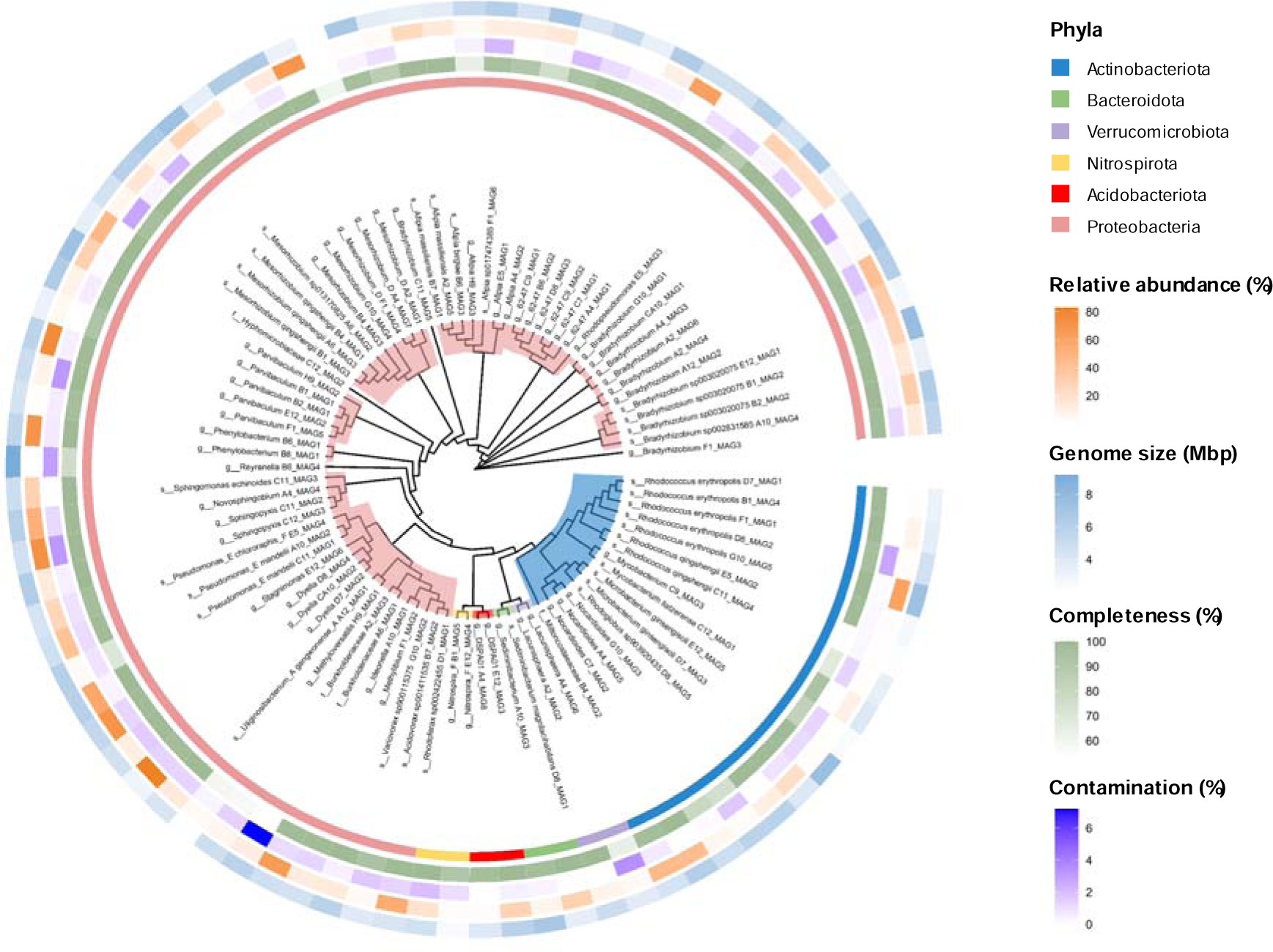
Phylogenetic tree of 88 MAGs derived from the 24 model communities based on 120 single-copy marker proteins for bacteria constructed using the maximum likelihood method. The taxonomy was classified by GTDB-Tk.

**Figure 5.**
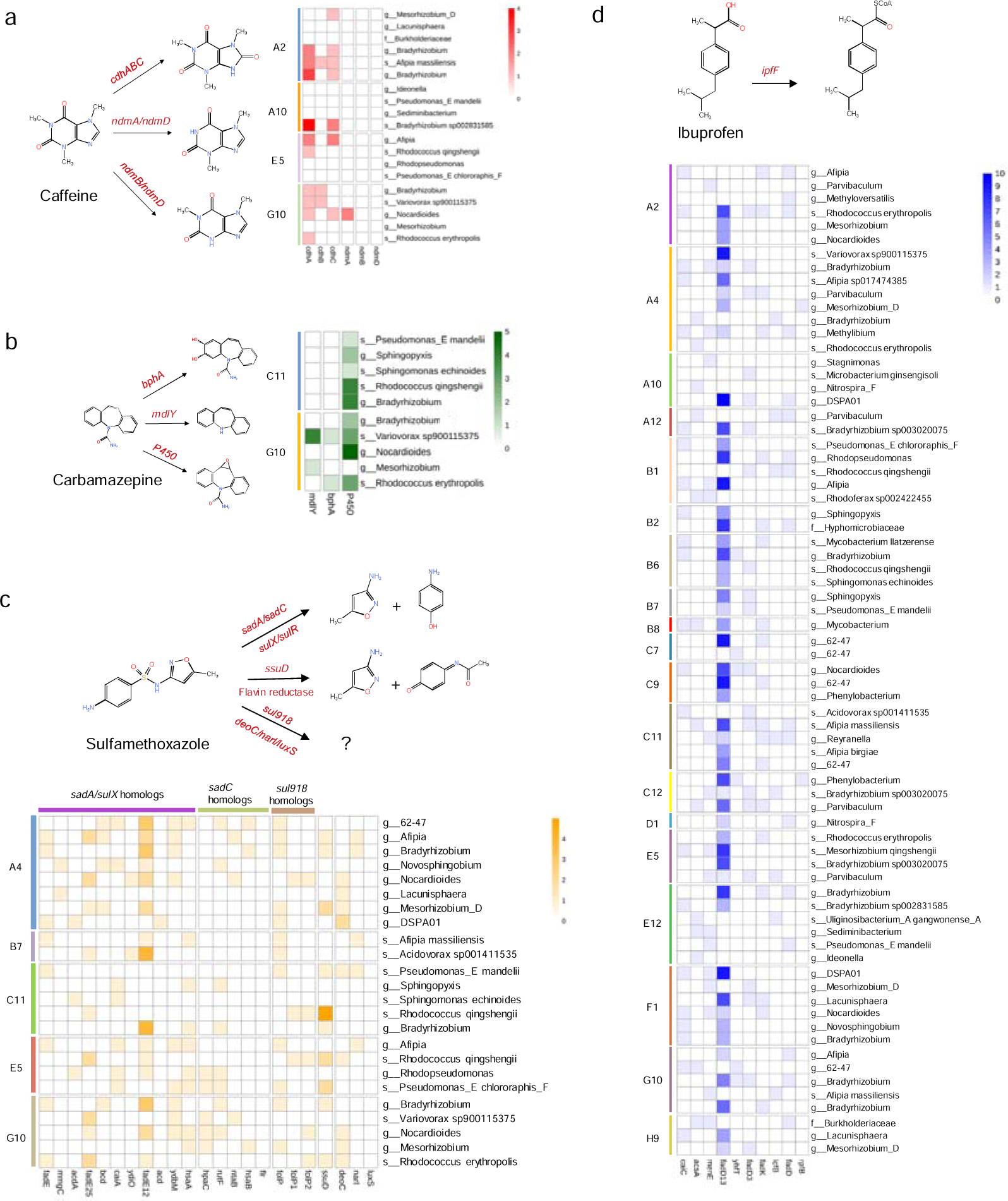
Initial biotransformation pathways, and the presence of related genes and their homologs in the degrading model communities of (a) caffeine, (b) carbamazepine, (c) sulfamethoxazole, and (d) ibuprofen. Values of the heatmap legend represent the number of genes identified in each MAG. Gabapentin was excluded since no biotransformation genes or enzymes have been reported yet. Trimethoprim was excluded since there was no degrading model community. Atenolol was not shown here but was described in the results section.

#### Caffeine

To date two distinct caffeine biotransformation pathways, *N*-demethylation and C-8 oxidation, has been uncovered in bacteria^18^. In the four caffeine-degrading model communities (A2, A10, E5 and G10), the biotransformation genes mainly appeared as the *cdhABC* genes (involved in C-8 oxidation) with only G10 harboring the *ndmA* gene (involved in *N*-demethylation) in one MAG classified to genus *Nocardiodes*. Since *ndmA* is highly dependent on *ndmD*, which is a partner reductase that transfers electrons to power the reaction, the incomplete pathway in G10 suggested the absence of caffeine *N*-demethylation. Hence, caffeine oxidation to 1,3,7-trimethyluric acid was the only pathway in all degrading model communities (Figure 5a). *Bradyrhizobium* carried the most abundant *cdh* genes in community A2, A10 and G10, and *Bradyrhizobium* sp002831585 was the only species carrying *cdhA* and *cdhC* in the community A10, indicating that *Bradyrhizobium* might have the potential of transforming caffeine. In the model community E5, *Afipia* carried *cdhA* and *cdhC* genes and species *Rhodococcus qingshengii* carried only *cdhA* gene. The complete removal of caffeine by E5 suggested that the novel species belonging to *Afipia* and the species *Rhodococcus qingshengii* might be the highly efficient caffeine degraders. Upon our survey of caffeine-degrading microorganisms, there have been no reports on caffeine biotransformation by *Bradyrhizobium*, *Afipia* and *Rhodococcus qingshengii* yet, but *Rhodococcus* sp. was found to degrade caffeine via oxidation in a mixture culture with *Klebsiella* sp.^79^. The subsequent transformation steps of 1,3,7-trimethyluric acid involved MAGs lacking *cdh* genes (Figure S2a), indicating the different roles of MAGs in assembling the caffeine oxidation pathway.

#### Carbamazepine

The first step of carbamazepine biotransformation was accomplished by BphA (biphenyl dioxygenase) metabolizing carbamazepine to cis-10,11-dihydroxy-10,11-dihydrocarbamazepine and cis-2,3-dihydroxy-2,3-dihydrocarbamazepine^80^, amide hydrolase removing the amide group^81^, and cytochrome P450 via monooxygenation to carbamazepine-10,11-epoxide^22^ (Figure 5b). Cytochrome P450 were present in all MAGs except for one MAG affiliated with *Mesorthizobium* of the two carbamazepine-degrading model communities C11 and G10. This is a ubiquitous enzyme system that is important for xenobiotic metabolism in bacteria catalyzing reactions such as aliphatic hydroxylations, epoxidations, and dealkylations^82^. Community C11 only contained cytochrome P450 with its most abundance observed in *Rhodococcus qingshengii* and *Bradyrhizobium*, indicating monooxygenation was the only biotransformation pathway in G10 and *Rhodococcus qingshengii* and *Bradyrhizobium* could play a critical role. The *bphA* gene and amide hydrolase encoding gene *mdlY* were present only in the community G10 with species *Variovorax* sp900115375 carrying the most abundant *mdlY*. *Variovorax* sp900115375 also contained *bphA*. *Mesorthizobium* that lacked the ubiquitous P450 harbored *mdlY*, and *Rhodococcus erythropolis* harbored *bphA*. The existence of *mdlY* and *bphA* genes in the model community G10 indicated that the metabolism of carbamazepine by G10 could undergo deamidation and hydroxylation pathways in addition to monooxygenation, which might explain the higher removal efficiency in G10 (40%) than C11 (23%). The deamidation could be attributed to *Variovorax* sp900115375 and *Mesorthizobium*, and the hydroxylation could be attributed to *Variovorax* sp900115375 and *Rhodococcus erythropolis*. Biotransformation of organic chemicals requires multifunctionality (multiple metabolic pathways), Stravs et al.^83^ suggested diverse biotransformation pathways supported by different detected byproducts could enhance the transformation of a broad range of micropollutants in freshwater phytoplankton, that is, functional (pathway) diversity benefits TOrC biotransformation. The unique presence of *bphB* for the next reaction step of carbamazepine hydroxylation in *Rhodococcus erythropolis* in G10 indicated the hydroxylation of carbamazepine was either shared by *Variovorax* sp900115375 and *Rhodococcus erythropolis*, or solely completed by *Rhodococcus erythropolis* (Figure S2b). This suggested the synergistic interaction between members in one model community contributes to TOrC metabolic processes, which were also observed in other recalcitrant chemical degradation studies^84,85,86^. Although the key degrading bacteria (usually responsible for the first transformation step) are important, the effective performance of a microbial community also depends on the populations targeting intermediates.

#### Sulfamethoxazole

The known pathways for sulfamethoxazole biotransformation are i) cleavage of the -C-S-N- bond in the sulfonamide molecules leading to 4-aminophenol and 3-amino-5-methylisoxazole by flavin dependent monooxygenase and reductase encoded by *sadA* and *sadC*, respectively^24^, ii) hydroxylation of aromatic ring by flavin monooxygenase encoded by *ssuD* and flavin reductase^87^. Liu et al.^88^ isolated a highly efficient sulfamethoxazole-degrading strain *Pseudomonas silesiensis* F6a. The *sadA* and *sadC* genes were not identified in its genome, and based on the detected metabolites several key functional genes (e.g., *deoC*, *narI*, *luxS*) participated in C-S cleavage, S-N hydrolysis and isoxazole ring cleavage were proposed. We identified the presence of homologs of SadA and SadC in the MAGs of five sulfamethoxazole-degrading model communities, only B7 did not contain any *sadC* homologs (Figure 5c). In principle, the function of *sadA* requires the assistance of *sadC*, thus the pathway of attacking the -C-S-N- bond was deactivated in B7. Moreover, community B7 did not carry *ssuD* gene, and only *folP* (homolog of sul918) and *narI* were present. The sulfonamides resistant gene *sul918* was reported to facilitate the removal of sulfamethoxazole, while itself cannot catalyze sulfamethoxazole. Hence, these observed results pointed out that *narI* could be the key biotransformation gene in B7 and *Afipia massiliensis* might be the potential sulfamethoxazole degrader. FadE12 (acyl-CoA dehydrogenase) was the most abundant SadA homologous protein which appeared mainly in *Bradyrhizobium*, *62-47* and *Acidovorax* sp001411535. *Rhodococcus qingshengii* in community C11 carried the dominant *ssuD* gene which is currently only reported on strain *Shewanella oneidensis* MR-1^87^, indicating its potential importance in sulfamethoxazole hydroxylation. The *sad* gene cluster was found to be conserved in two genera *Paenarthrobacter* and *Microbacterium*^71^, it is consistent in our study that no *sadA* or *sadC* was observed in the 88 MAGs with diverse taxonomy. Nevertheless, our study suggested that the functions of *sad* gene cluster could be taken over by their homologs.

#### Ibuprofen

The *ipfF* gene encoding ibuprofen ligase is responsible for transforming ibuprofen to ibuprofen-CoA^89^. Although *ipfF* was not identified in the degrading model communities, its homologs were characterized (Figure 3a). All 72 MAGs of degrading communities harbored at least one of the IpfF homologous proteins with *fadD13* exhibiting the highest abundance in genera *Bradyrhizobium*, *62-47*, *Rhodococcus*, *Sphingopyxis*, *Mycobacterium* and *Rhodopseudomonas* (Figure 5d). The biodiversity of ibuprofen-degrading model communities revealed by species richness varied from one to eight at the MAG level. Stadler et al.^90^ established cultures with a gradient of microbial biodiversity from activated sludge via dilution-to-extinction, and found the loss of biodiversity had a significant correlation with the reduction of biotransformation for atenolol, carbamazepine and venlafaxine. However, ibuprofen biotransformation degree was not affected by the species richness with all degrading communities exhibiting 100% removal. Notably, community B8 and D1 consisted of only single MAG affiliated with *Phenylobacterium* and *Rhodoferax* sp002422455, respectively, indicating their unique function in degrading ibuprofen. These two bacteria were reported for the first time in ibuprofen efficient biotransformation. More interestingly, D1 contained only *fadD* and *fadD13* encoding enzyme long-chain-fatty-acid-CoA ligase. This indicated that the long-chain-fatty-acid-CoA ligase is the critical enzyme in ibuprofen biotransformation, and the *fadD* or *fadD13* gene could take the place of *ipfF* gene. The wide distribution of *fadD* and *fadD13* genes also suggested that the CoA ligation to ibuprofen could be driven by diverse bacteria. The existing of other homologs and aliphatic monooxygenase (present in all communities) need further research to confirm whether and how they play a part, which could be supported by transformation products determination, biotransformation experiment on extracted enzyme, and transcripts indicating gene expression.

#### Atenolol and gabapentin

The first step of atenolol biotransformation pathway was reported to be the acetylation of the amino group catalyzed by amidohydrolase, and the related bacterium was *Hydrogenophaga*^91^. In the present study, amidohydrolase was identified in all 88 MAGs, while only three communities can degrade atenolol. Ammonia monooxygenase was also found to convert atenolol to atenolol acid by ammonia oxidizing bacteria^92^, and it only appeared in *Dyella* in D8 community and was absent in the other two atenolol-degrading community C7 and C9. The roles of amidohydrolase and ammonia monooxygenase still need further investigation regarding their expression or activity. Gabapentin was reported to be transformed to gabapentin-lactam via intramolecular amidation in the biological process^93^ while the related enzymes have not been addressed yet. Community B4 and F1 removed 32% and 42% of gabapentin, respectively. Our results indicated that there could also be novel functions and pathways for atenolol and gabapentin biotransformation that are unmined.

In summary, in this section we discussed the known biotransformation genes, enzymes and pathways for caffeine, carbamazepine, ibuprofen, sulfamethoxazole, atenolol and gabapentin. Trimethorpim was excluded since no model community showed removal on it. We then related the distribution of these biotransformation agents to the removal efficiencies of model communities, and inferred potential associated degrading bacteria and functional alternatives (homologs). We found that caffeine oxidation to 1,3,7-trimethyluric acid was the dominant pathway, with *Bradyrhizobium*, *Afipia* and *Rhodococcus qingshengii* acting as potential degraders that are reported for the first time. Carbamazepine biotransformation could be enhanced by the involvement of cytochrome P450, *mdlY* and *bphA* providing multiple pathways, *Rhodococcus qingshengii*, *Rhodococcus erythropolis*, *Bradyrhizobium, Variovorax sp900115375 and Mesorthizobium* might be the associated bacteria. The long-chain-fatty-acid-CoA ligase was found to be the critical enzyme in ibuprofen biotransformation, and the *fadD* and *fadD13* gene could function as *ipfF* gene. Efficient ibuprofen biotransformation could be driven by diverse microorganisms. The *sad* gene cluster responsible for sulfamethoxazole biotransformation was not identified in the degrading model communities, but *sadA*’s homolog *fadE12* was abundant in *Bradyrhizobium*, *62-47* and *Acidovorax sp001411535*. Notably, *Rhodococcus qingshengii* has been reported to be able to degrade carbendazim^94^, phenanthrene^95^, triphenylmethane dyes^96^, phenol^97^, naphthalene^98^, and crude oil^99^. Our results showed that *Rhodococcus qingshengii* also carried caffeine, carbamazepine, sulfamethoxazole and ibuprofen biotransformation genes and enzymes, and the removal performance indicated its potential ability in various TOrC metabolism. For atenolol, gabapentin and trimethoprim, since their current biotransformation knowledge is scarce, more experiments on degrading model communities (e.g., pure culture isolation, transformation products detection, enzyme-based degradation, gene expression) are required to confirm the function of reported enzymes (e.g., amidase, amidohydrolase) and to explore novel pathways.

### Comparative genomic analyses indicated potential novel functions

The known pathways and biotransformation agents discussed in the above sections provided us with an overview of their existence in the 24 model communities. However, the unknown mechanisms still need further research to deepen our understanding of various TOrC biotransformation by diverse species. Here, we conducted comparative genomic analyses via GO enrichment and related gene family prediction, aiming to discover potential novel functions shared by model communities with similar TOrC removal performance. The GO enrichment analysis compared the significantly enriched functions among the 24 model communities (Figure 6a). In general, most model communities possessed their unique functions, while B4 and A6, C9 and C12 shared similar GO modules. To be specific, community A6 consisted of a dominant strain belonging to Burkholderiaceae (69.8%) and two *Mesorhizobium* strains (13.8% and 0.7%), B4 was dominated by *Mesorhizobium* (75.9% and 8.3%) and 4.3% was from Miltoncostaeales. Although they were dominated by different bacteria, their biological functions were both enriched in sarcosine oxidase activity, tetrahydrofolate metabolic process, methanogenesis, polyamine transmembrane transport and carbon-sulfur lyase activity, and these two communities showed similar TOrC removal performance that can only biotransform gabapentin (A6: 21% and B4: 32%). The community C9 which could biotransform 100% ibuprofen and atenolol was composed by three strains belonging to genera *62-47* (26.8% and 40.4%) and *Mycobacterium* (3.4%), and community C12 degrading only ibuprofen consisted of three strains from species *Mycobacterium llatzerense* (13.2%), genus *Sphingopyxis* (54.7%) and family Hyphomicroblaceae (3.8%). Peptidoglycan-based cell wall, regulation of cellular biosynthetic process, glucose-6-phosphate dehydrogenase (coenzyme F420) activity, fatty-acyl-CoA synthase activity, and response to abiotic stimulus were the functions shared by C9 and C12. The community E5 was the best degrader that biotransformed the most TOrCs (i.e., caffeine, ibuprofen, sulfamethoxazole, carbamazepine and gabapentin), showing the unique enrichment in glutamate synthase (NADPH) activity, bacteriochlorophyll binding, toxic substance binding and plasma membrane light-harvesting complex. The GO enriched analysis in model communities could indicate the association between these enriched functions and specific TOrC biotransformation. A supportive example is that the fatty-acyl-CoA synthase activity enriched in ibuprofen-degrading communities (C9 and C12) is related to transformation of ibuprofen to ibuprofen-CoA.

**Figure 6.**
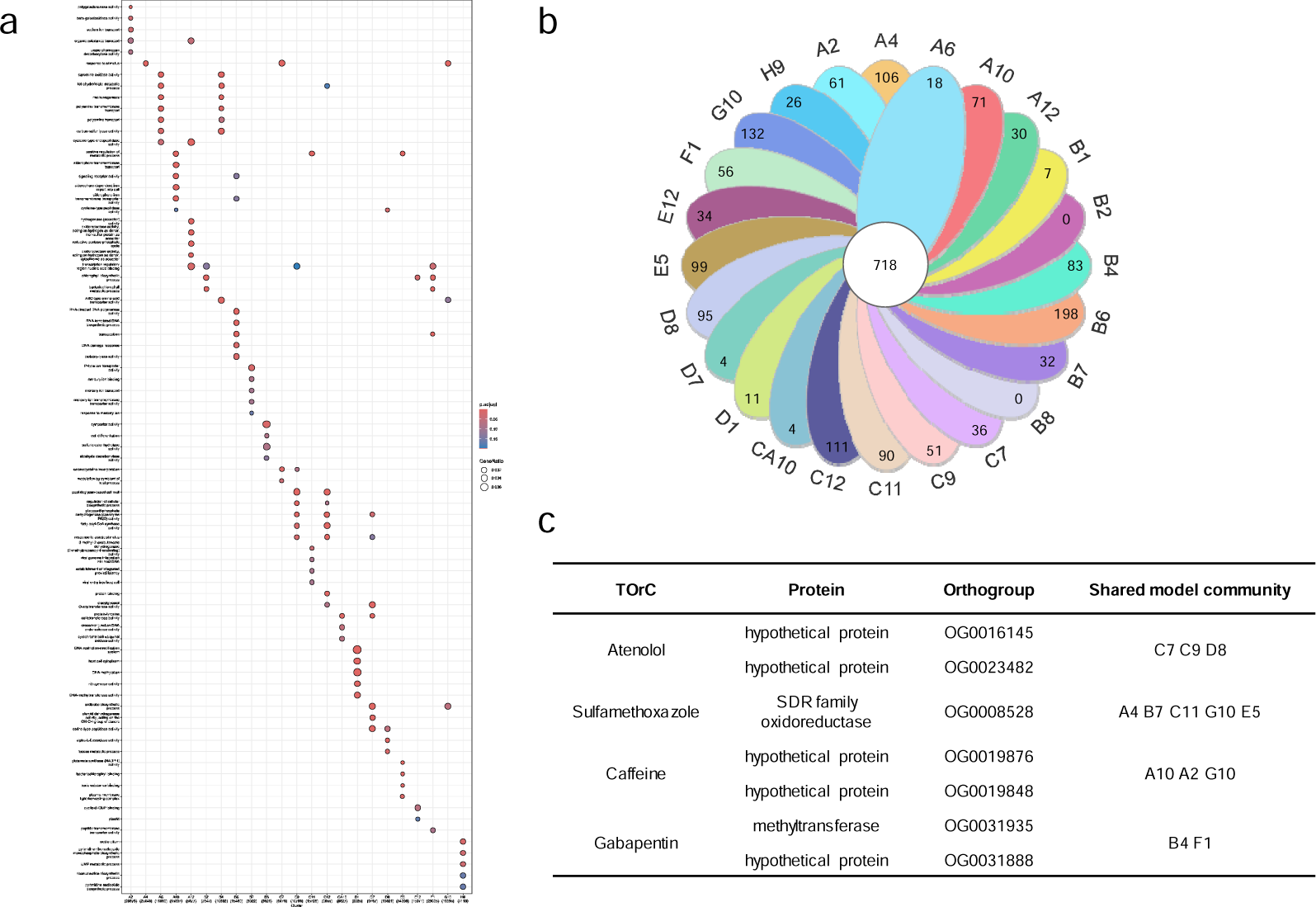
Comparative genomic analyses unraveling potential uncharacterized biotransformation functions. (a) Gene ontology (GO) functional annotation and enrichment of the 24 model communities. The x axis is model communities with the number of genes annotated with GO terms. The y axis is the description of GO terms. The color and size of the dots represent the significance of GO terms, and the gene ratio (the percentage of differentially expressed genes in a given GO term), respectively. (b) Venn diagram showing the shared and unique orthogroups among the 24 model communities identified using OrthoFinder. (c) Shared orthogroups of TOrC specific degrading model communities indicating putative novel biotransformation enzymes. The sequences of each orthogroup were provided in Table S5.

Furthermore, in order to find uncharacterized putative biotransformation functions, we used OrthoFinder to access orthologous clusters of the 24 model communities. As a result, OrthoFinder assigned 414772 genes (94.5 % of total) to 36064 orthogroups, 718 of which were found in all 24 concatenated genomes, and 1355 of which were community-specific orthogroups that were only present in one genome (Figure 6b). The sulfamethoxazole degrader A4, B7, C11, G10 and E5 shared only one orthogroup with the function of SDR (short-chain dehydrogenases/reductases) family oxidoreductase. SDR enzymes play important roles in lipid, amino acid, carbohydrate, cofactor, hormone and xenobiotic metabolism^100^. Interestingly, some studies have shown that the SDR family enzymes are upregulated in microbes when they are challenged with organic pollutants^101,102,103,104^. These findings support our preliminary data that the SDR family oxidoreductase could be involved in sulfamethoxazole biotransformation. There were two hypothetical protein orthogroups identified in caffeine degrading community A10, A2 and G10 and no shared orthogroups were found when we included E5. This might suggest that the difference of caffeine removal efficiency between E5 and the other three communities (A10, A2, G10) might be attributed to their functional divergence. The atenolol degrader C7, C9 and D8 shared two orthogroups identified as hypothetical protein that have not been annotated yet. For gabapentin degrading community B4 and F1, they shared one hypothetical protein orthogroup and another characterized as methyltransferase. Considering the chemical structure of gabapentin, it is unlikely the methyltransferase would act on the functional groups or ring in the first step of transformation, but it might be involved in subsequent reactions if gabapentin is attacked by other enzymes resulting in structural modification. Nevertheless, our inference needs further experiments to validate. Moreover, the functional annotation of identified promising hypothetical proteins (sequences were provided in Table S6) also requires further investigation in our future research.

Hence, by using the comparative analyses we proposed enriched functions and putative biotransformation enzymes based on the TOrC-degrading and non-degrading model communities (Table S6). However, since the model communities were shaped by diverse species and showed different degrees of TOrC removal, the prediction of gene families related to each TOrC biotransformation is not that straightforward. In the future work, it could be more reliable to apply transcriptomic analysis to analyze the upregulated enzymes or differential expressed genes.

## Conclusions

In this study, we obtained 24 bacterial model communities with one to eight taxa by adapting them to seven TOrCs (i.e., atenolol, caffeine, carbamazepine, gabapentin, ibuprofen, sulfamethoxazole, trimethoprim) prior to dilution-to-extinction. In addition, we profiled the biotransformation genes, enzymes and associated bacteria of each TOrC by metagenome-centric analyses integrated with currently known biotransformation knowledge. Our research was conducted on adaptation-dilution-cultivation model communities in response to real-world TOrC concentrations, filling out the knowledge for both well-understood chemicals (e.g., caffeine) and less well-understood TOrCs (e.g., carbamazepine, gabapentin). Our main findings are:

1. The 24 model communities exhibited different TOrC removal abilities and we achieved several efficient degraders for ibuprofen (100% removal), caffeine (100% removal) and atenolol (85-100% removal). The transformation efficiencies for other TOrCs ranged from 0% to 45% with almost no removal on trimethoprim. The community E5 was the best degrader with the ability of biotransforming multiple organic chemicals of diverse structures: caffeine, ibuprofen, sulfamethoxazole, carbamazepine and gabapentin.
2. The presence of initial biotransformation genes and enzymes did not fully support the corresponding TOrC removal, and further expression level validation is needed. Functional similar homologs to existing biotransformation genes and enzymes could play critical roles in TOrC metabolism. Long-chain-fatty-acid-CoA ligase encoded by *fadD* and *fadD13* gene could be responsible for CoA ligation to ibuprofen. Acyl-CoA dehydrogenase encoded by *fadE12* gene could function as SadA to transform sulfamethoxazole by attacking the -C-S-N-bond.
3. Novel TOrC-degraders were reported for the first time. *Bradyrhizobium*, *Afipia* and *Rhodococcus qingshengii* were potential caffeine-degrading bacteria. *Rhodococcus qingshengii*, *Rhodococcus erythropolis*, *Bradyrhizobium, Variovorax sp900115375 and Mesorthizobium* might be carbamazepine-degrading associated bacteria. *Bradyrhizobium*, *62-47* and *Acidovorax sp001411535* could be responsible for sulfamethoxazole biotransformation. *Rhodococcus qingshengii* carrying caffeine, carbamazepine, sulfamethoxazole and ibuprofen biotransformation genes and enzymes could be a promising species for multiple TOrC removal.
4. SDR family oxidoreductase could be involved in sulfamethoxazole biotransformation. Novel putative hypothetical proteins were identified in caffeine, atenolol and gabapentin degrading model communities, but their functions as well as resulting pathways require further analysis.

## Data availability

Raw metagenome sequencing data and assembled MAGs have been deposited at INSDC (with ENA: https://www.ebi.ac.uk/ena) under the project accession number PRJEB74141.

## Acknowledgments

This study was funded by the German Research Foundation (DFG, WU 890/2-1) and China Scholarship Council (LC). We are grateful to Leibniz-Rechenzentrum for providing computational resources. Ignacio Fernando Sottorff Neculhueque and Javad Ahmadi are thanked for assisting the LC-MS/MS measurement. Laboratory for Functional Genome Analysis (LAFUGA) of the Ludwig-Maximilian-University Munich is thanked for performing the high-throughput sequencing.

## Author contributions

LC and CW designed this research; LC performed the experiments, analyzed the data and drafted the manuscript; CW and SLG guided the data analysis and revised the manuscript. All authors read and approved the final manuscript.

