## Supplementary Information for "Profiling trace organic chemical biotransformation genes, enzymes and associated bacteria in microbial model communities"

Table S1. The names, structure, uses, occurrence, RQ values and biotransformation efficiencies of 27 TOrCs used in this study.

| Compound | Structure | Use | Occurrence (μg/L) | RQ | Biotransformation efficiency (%) | Reference |
|---|---|---|---|---|---|---|
| Atenolol | 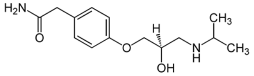 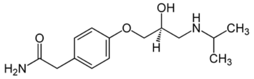 | Beta-Blockers | ND-26.5 | 0-0.014 | <0-85.1 | [1-3] |
| Caffeine | 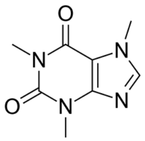 | Psychoactive drug | ND-220 | 0.02 | 49.9-99.6 | [1] [2] [4] |
| Carbamazepine | 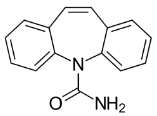 | Antiepileptics | ND-38.24 | 0-1.8 | <0-62.3 | [1] [2] |
| Gabapentin | 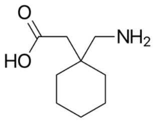 | Anticonvulsant drug | 0.79-15.36 | 1.32 | 7.9 | [2] [5] [6] |
| Ibuprofen | 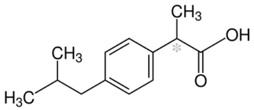 | Antirheumatic drug | ND-303.0 | 0-5.42 | 72-100 | [1] [2] [7] |
| Sulfamethoxazole | 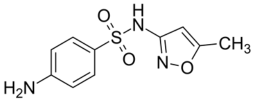 | Antibacterials | ND-54.8 | 0-74.4 | 4-88.9 | [1] [2] [4] |
| Trimethoprim | 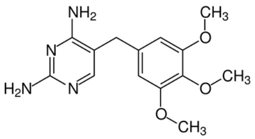 | Antibacterials | 13.6 | 2.3 | <0-81.6 | [1] [2] [8] |

Table S2. Treatment conditions of the enrichment experiments and the preliminary selection of model communities.

| TOrC combination for enrichment | Number of diluted communities | Number of positive growth communities after dilution | Number of communities after TOrC removal measurement | Number of communities after taxonomy refinement |
|---|---|---|---|---|
| Atenolol (ATN) | 96 | 12 | 10 | 2 |
| Caffeine (CAF) | 96 | 5 | 4 | 2 |
| Ibuprofen (IBU) | 96 | 16 | 16 | 4 |
| Sulfamethoxazol (SMX) | 96 | 20 | 8 | 2 |
| Trimethoprim (TMP) | 96 | 6 | 0 | 0 |
| Gabapentin (GAP) | 96 | 9 | 1 | 1 |
| Carbamazepine (CBZ) | 96 | 11 | 3 | 1 |
| CBZ + GAP + TMP | 96 | 4 | 3 | 2 |
| CAF + IBU + SMX + ATN | 96 | 9 | 5 | 2 |
| CBZ + CAF + IBU + SMX + ATN | 96 | 15 | 2 | 1 |
| CBZ + TMP + GAP + CAF + IBU + SMX + ATN | 96 | 14 | 2 | 1 |

Table S3. Currently known biotransformation genes, enzymes, pathways, and associated bacteria of seven TOrCs investigated in this study.

| **Compound** | **Gene** | **Protein** | **Strain** | **Pathway** | **Reference** |
|---|---|---|---|---|---|
| Caffeine | *cdhABC* | Caffeine dehydrogenase | *Pseudomonas putida* CBB1<br>*Klebsiella*<br>*Rhodococcus*<br>*Alcaligenes* sp. | Caffeine -----> 1,3,7-Trimethyluric acid | [9-11] |
|  | *tmuM* | FAD-binding monooxygenase |  | 1,3,7-Trimethyluric acid ----> 1,3,7-Trimethyl-5-hydroxyisouric acid |  |
|  | *tmuH* | 5-hydroxyisourate hydrolase |  | 1,3,7-Trimethyl-5-hydroxyisouric acid ----> 3,6,8-Trimethyl-2-oxo-4-hydroxy-4-carboxy-5-ureidoimidazoline |  |
|  | *tmuD* | 2-oxo-4-hydroxy-4-carboxy-5-ureidoimidazoline decarboxylase |  | 3,6,8-Trimethyl-2-oxo-4-hydroxy-4-carboxy-5-ureidoimidazoline ----> S-(+)-3,6,8-trimethyl-allantoin |  |
|  | *ndmA* | Methylxanthine N1-demethylase | *Pseudomonas putida* CBB5<br>*Pseudomonas putida*<br>*Pseudomonas* sp.<br>*Serratia marcescens*<br>*Paraburkholderia caffeinilytica* CF1<br>*Brevibacterium* sp.<br>*Leifsonia* sp. SIU | Caffeine -----> Theobromine | [12-16] |
|  | *ndmB* | Methylxanthine N3-demethylase |  | Theobromine -----> 7-Methyxanthine<br>Caffeine -----> Paraxanthine |  |
|  | *ndmC* | Methylxanthine N7-demethylase |  | 7-Methyxanthine -----> Xanthine |  |
| | *ndmD* | Reductase | | / | |
| | *ndmE* | Glutathione S-transferase | | / | |
| Ibuprofen | *ipfF* | Long chain fatty acid--CoA ligase | *Sphigomonas* sp. Ibu-2<br>*Rhizorhabdus wittichii* MPO218<br>*Sphingopyxis granuli* RW412<br>*Pseudomonas citronellolis* | Ibuprofen -----> Ibuprofen-CoA | [17-20] |
|  | *ipfABHI* | Aromatic ring dioxygenase |  | Ibuprofen-CoA -----> Dihydroxyibuprofen-CoA |  |
|  | *ipfDE* | Thiolase<br>OB-fold domain-containing protein |  | Dihydroxyibuprofen-CoA -----> Isobutylcatechol<br>Dihydroxyibuprofen-CoA -----> Propionyl-CoA |  |
|  | *ipfL* | Catechol-2,3-dioxygenase |  | Isobutylcatechol -----> 5-Isobutyl-2-hydroxymuconate semialdehyde |  |
|  | *ipfM* | 5-carboxymethyl-2-hydroxymyconic semialdehyde dehydrogenase |  | 5-Isobutyl-2-hydroxymuconate semialdehyde -----> 5-Isobutyl-2-hydroxymuconic acid |  |
|  | *pccAB* | Propionyl-CoA carboxylase |  | Propionyl-CoA -----> S-methylmalonyl-CoA |  |
|  | / | Meta ring fission enzymes | *Variovorax* Ibu-1 | Ibuprofen -----> Trihydroxyibuprofen | [21] |
|  | / | / | *Nocardia* sp. NRRL 5646 | Ibuprofen -----> Ibuprophenol | [22] |
|  | / | Aliphatic monooxygenase | *Pseudoalteromonas* sp. | Ibuprofen -----> 2-Hydroxyibuprofen | [23-25] |

| | | | | | |
|---|---|---|---|---|---|
| | | | *Citrobacter freundii* PYI-2<br>*Citrobacter portucalensis* YPI-2<br>*Bacillus thuringiensis* B1 | | |
|  | / | Synthase acyl-CoA | *Citrobacter freundii* PYI-2<br>*Citrobacter portucalensis* YPI-2 | 2-(4-Hydroxyphenyl-) propionic acid -----> 1,4-Hydroquinone | [24] |
|  | / | Hydroquinone monooxygenase |  | 1,4-Hydroquinone -----> 2-Hydroxy-1,4-quinol |  |
|  | / | Hydroxyquinol 1,2-dioxygenase |  | 2-Hydroxy-1,4-quinol -----> 3-Hydroxy-cis,cis-muconic acid |  |
| Sulfamethoxazole | *sadA* | FMNH2-dependent monooxygenases | *Microbacterium* sp. BR1/C448<br>*Actinobacteria* sp.<br>*Paracoccus denitrificans* DYTN-1<br>*Paenarthrobacter* sp. P27<br>*Paenarthrobacter ureafaciens* YL1 | Sulfamethoxazole -----> 4-Aminophenol + 3-Amino-5-methylisoxazole | [26-31] |
|  | *sadB* | FMNH2-dependent monooxygenases |  | 4-Benzoquinone imine -----> 1,2,4-Trihydroxybenzene |  |
| | *sadC* | Flavin reductase | | / | |
|  | *ssuD* | Flavin monooxygenase | *S. oneidensis* MR-1 | Sulfamethoxazole -----> N-acetyl-p-benzoquinoneimine + 3-Amino-5-methylisoxazole | [32] |
|  | *sulX* | sulfonamide monooxygenase | Microbacterium sp. CJ77 | Sulfamethoxazole -----> 4-Aminophenol + 3-Amino-5-methylisoxazole | [33] |
| | *sulR* | flavin reductase | | / | |
| | *sul918* | Sulfonamide-resistant dihydropteroate synthase | *Paenarthrobacter* sp. SD-1 | / | [34] |
|  | / | Group D flavin monooxygenase | *Achromobacter denitrificans* PR1 and *Leucobacter* sp. GP | Sulfamethoxazole -----> *Ipso*-substituted sulfamethoxazole | [35] |
| | *deoC, narI, luxS, nuoH, unknown Gene0655, unknown Gene4650* | / | *Pseudomonas silesiensis* F6a | C-S cleavage, S-N hydrolysis, Isoxazole ring cleavage | [36] |
| Atenolol | / | Amidohydrolase | *Hydrogenophaga* sp. YM1 | Atenolol -----> Atenolol acid | [3] |
|  | *tfdA* | 2,4-dichlorophenoxyacetate dioxygenase |  | Atenolol acid -----> 4-Hydroxyphenylacetic acid |  |

| | | | | | |
|---|---|---|---|---|---|
|  | / | Ammonia monooxygenase | Ammonia oxidizing bacteria | Atenolol -----> Atenolol acid | [37] |
| Carbamazepine | *bphA* | Biphenyl dioxygenases | *Paraburkholderia xenovorans* LB400 | Carbamazepine -----> C*is*-10,11-dihydroxy-10,11-dihydrocarbamazepine + C*is*-2,3-dihydroxy-2,3-dihydrocarbamazepine | [38] |
|  | *bphB* | Hydrodiol dehydrogenase |  | *Cis*-2,3-dihydroxy-2,3-dihydrocarbamazepine -----> 2,3-Dihydroxycarbamazepine |  |
|  | / | Amide hydrolase | *Gordonia polyophrenivorans* | Carbamazepine -----> Iminostilbene | [39] |
|  | / | Dioxygenase |  | Iminostilbene -----> Dibenzazoheteroketone |  |
|  | / | Lyase and aldolase |  | Dibenzazoheteroketone -----> Diphenylamine |  |
|  | / | Cytochrome P450 | *Labrys portucalensis* F11 | Carbamazepine -----> Carbamazepine-10,11-epoxide + Iminostilbene | [40] |
| Trimethoprim | / | Ammonia monooxygenase | Ammonia oxidizing bacteria | / | [41] |
|  | / | Demethylase | / | Trimethoprim -----> 4-desmethyl-trimethoprim | [42] |
| Gabapentin | / | / | *Micrococcus luteus* N.ISM.1 | / | [43] |
|  | / | / | / | Gabapentin -----> Gabapentin-lactam | [44] |

Table S4. Initial biotransformation genes and their homologs identified using OrthoFind.

| Gene | Number of homologs | Domain | Molecular function | Accession number (UniProtKB) |
|---|---|---|---|---|
| *cdhA* | 27 | Ald_Xan_dh_C<br>Ald_Xan_dh_C2 (70.37%) | oxidoreductase activity (74.07%) | D7REY3 |
| *cdhB* | 23 | CO_deh_flav_C<br>FAD_binding_5 (60.87%) | flavin adenine dinucleotide binding (91.30%)<br>UDP-N-acetylmuramate dehydrogenase activity (82.61%) | D7REY4 |
| *cdhC* | 18 | Fer2<br>Fer2_2 (94.44%) | electron carrier activity (100.00%)<br>metal ion binding (100.00%)<br>2 iron, 2 sulfur cluster binding (94.44%)<br>oxidoreductase activity (88.89%) | D7REY5 |
| *ndmA* | 10 | Rieske (70.00%) | 2 iron, 2 sulfur cluster binding (100.00%)<br>metal ion binding (50.00%) | H9N289 |
| *ndmB* | 8 | Rieske (87.50%) | 2 iron, 2 sulfur cluster binding (100.00%)<br>metal ion binding (62.50%)<br>oxidoreductase activity (50.00%) | H9N290 |
| *ndmD* | 83 | FAD_binding_6<br>Fer2<br>NAD_binding_1 (38.55%) | 2 iron, 2 sulfur cluster binding (69.88%)<br>metal ion binding (60.24%)<br>electron carrier activity (60.24%)<br>oxidoreductase activity (54.22%) | H9N291 |
| *ipfF* | 59 | AMP-binding<br>AMP-binding_C (98.31%) | ligase activity (37.29%) | A1E027 |
| *sadA* | 82 | Acyl-CoA_dh_1<br>Acyl-CoA_dh_M<br>Acyl-CoA_dh_N (73.17%) | flavin adenine dinucleotide binding (98.78%)<br>acyl-CoA dehydrogenase activity (98.78%) | A0A3G2JSU3 |
| *sadC* | 47 | Flavin_Reduct (97.87%) | riboflavin reductase (NADPH) activity (97.87%)<br>FMN binding (97.87%) | / |

| | | | | |
|---|---|---|---|---|
| *sulX* | 106 | Acyl-CoA_dh_1<br>Acyl-CoA_dh_M<br>Acyl-CoA_dh_N (68.87%) | flavin adenine dinucleotide binding (98.11%)<br>acyl-CoA dehydrogenase activity (98.11%) | A0A482P9Z9 |
| *sulR* | 47 | Flavin_Reduct (97.87%) | riboflavin reductase (NADPH) activity (97.87%)<br>FMN binding (97.87%) | / |
| *ssuD* | 99 | Bac_luciferase (100.00%) | alkanesulfonate monooxygenase activity (100.00%) | Q9HYG2 |
| *sul918* | 27 | Pterin_bind (96.30%) | dihydropteroate synthase activity (95.92%) | / |
| *deoC* | 21 | Pentose phosphate pathway (98.92%) | deoxyribose-phosphate aldolase activity (100.00%) | P0A6L0 |
| *narI* | 14 | Nitrate_red_gam (100.00%) | nitrate reductase activity (100.00%) | P42177 |
| *luxS* | 15 | LuxS (100.00%) | iron ion binding (100.00%)<br>S-ribosylhomocysteine lyase activity (100.00%) | O34667 |
| *mdlY* | 41 | Amidase (100.00%) | carbon-nitrogen ligase activity, with glutamine as amido-N-donor (60.98%) | Q84DC4 |
| *bphA* | 42 | Rieske (52.38%) | 2 iron, 2 sulfur cluster binding (100.00%) | Q46372 |

Table S5. Completeness, contamination and relative abundance of 88 MAGs derived from the 24 model communities.

| MAG | Completeness (%) | Contamination (%) | Abundance (%) |
|---|---|---|---|
| A10_MAG_1 | 98.48 | 1.2 | 13.2 |
| A10_MAG_2 | 99.93 | 0.4 | 32 |
| A10_MAG_3 | 99.1 | 0.33 | 15.4 |
| A10_MAG_4 | 99.48 | 0.37 | 28 |
| A12_MAG_1 | 98.34 | 0.83 | 32.7 |
| A12_MAG_2 | 97.99 | 1.82 | 42 |
| A2_MAG_1 | 96.57 | 0.36 | 2.5 |
| A2_MAG_2 | 98.65 | 2.7 | 1.8 |
| A2_MAG_3 | 99.53 | 0 | 21.4 |
| A2_MAG_4 | 98.41 | 1.41 | 22.5 |
| A2_MAG_5 | 100 | 0.32 | 28.7 |
| A2_MAG_6 | 88.07 | 1.49 | 0.7 |
| A4_MAG_1 | 98.24 | 0.93 | 2.3 |
| A4_MAG_2 | 100 | 0 | 62 |
| A4_MAG_3 | 99.35 | 0.39 | 10.9 |
| A4_MAG_4 | 97.03 | 0.51 | 0.7 |
| A4_MAG_5 | 98.27 | 1.04 | 0.9 |
| A4_MAG_6 | 93.65 | 2.03 | 0.7 |
| A4_MAG_7 | 96.41 | 0.36 | 2.3 |
| A4_MAG_8 | 60.11 | 0 | 0.2 |
| A6_MAG_1 | 99.53 | 0 | 69.8 |
| A6_MAG_2 | 99.51 | 0.7 | 13.8 |
| A6_MAG_3 | 97.87 | 0.2 | 0.7 |
| B1_MAG_1 | 100 | 0 | 16.7 |
| B1_MAG_2 | 99.96 | 0.13 | 27.7 |
| B1_MAG_3 | 99.23 | 2.23 | 7 |
| B1_MAG_4 | 99.94 | 0.38 | 16.3 |
| B1_MAG_5 | 96.76 | 2.78 | 6.1 |
| B2_MAG_1 | 100 | 0 | 46.3 |

| | | | |
|---|---|---|---|
| B2_MAG_2 | 99.96 | 0.13 | 30 |
| B4_MAG_1 | 97.98 | 0.55 | 8.3 |
| B4_MAG_2 | 92.82 | 0 | 4.3 |
| B4_MAG_3 | 99.51 | 0.49 | 75.9 |
| B6_MAG_1 | 99.51 | 3.19 | 7.8 |
| B6_MAG_2 | 96.96 | 0.56 | 0.8 |
| B6_MAG_3 | 99.97 | 0.63 | 70.2 |
| B6_MAG_4 | 76.73 | 2.99 | 0.2 |
| B7_MAG__1 | 99.68 | 0 | 29.4 |
| B7_MAG__2 | 99.62 | 1.16 | 51.7 |
| B8_MAG_1 | 99.51 | 3.19 | 72.9 |
| C11_MAG_1 | 99.77 | 0.48 | 13.8 |
| C11_MAG_2 | 99.5 | 1.28 | 13.8 |
| C11_MAG_3 | 59.12 | 0.34 | 3.4 |
| C11_MAG_4 | 57.44 | 1.38 | 5.9 |
| C11_MAG_5 | 53.22 | 7.22 | 11.6 |
| C12_MAG_1 | 99.75 | 0.44 | 13.2 |
| C12_MAG_2 | 94.07 | 1.65 | 3.8 |
| C12_MAG_3 | 99.52 | 1.28 | 54.7 |
| C7_MAG_1 | 98.29 | 1.11 | 65.8 |
| C7_MAG_2 | 98.45 | 1.3 | 12.5 |
| C9_MAG_1 | 97.11 | 0.9 | 26.8 |
| C9_MAG_2 | 98.29 | 1.11 | 40.4 |
| C9_MAG_3 | 69.59 | 0.84 | 3.4 |
| CA10_MAG_1 | 98.91 | 1.25 | 70.5 |
| CA10_MAG_2 | 99.59 | 1.56 | 10.6 |
| D1_MAG_1 | 98.07 | 1.33 | 82.8 |
| D7_MAG_1 | 99.94 | 0.38 | 57.8 |
| D7_MAG_2 | 99.66 | 1.97 | 18.5 |
| D7_MAG_3 | 91.8 | 1.01 | 3 |
| D8_MAG_1 | 97.29 | 0 | 2.5 |

| | | | |
|---|---|---|---|
| D8_MAG_2 | 99.75 | 0.42 | 29.1 |
| D8_MAG_3 | 97.42 | 0.63 | 45.6 |
| D8_MAG_4 | 67.24 | 1.33 | 0.8 |
| D8_MAG_5 | 53.97 | 0.25 | 0.8 |
| E12_MAG_1 | 99.96 | 0.13 | 60.6 |
| E12_MAG_2 | 100 | 0 | 7.3 |
| E12_MAG_3 | 99.57 | 0 | 1.3 |
| E12_MAG_4 | 96.76 | 2.78 | 3 |
| E12_MAG_5 | 98.89 | 0.25 | 1.9 |
| E12_MAG_6 | 83.66 | 1.29 | 0.5 |
| E5_MAG_1 | 93.95 | 0 | 9.1 |
| E5_MAG_2 | 99.94 | 0 | 43.8 |
| E5_MAG_3 | 79.22 | 0.11 | 6.1 |
| E5_MAG_4 | 74.35 | 1.71 | 3 |
| F1_MAG_1 | 99.75 | 0 | 11.1 |
| F1_MAG_2 | 99.38 | 0.86 | 20.9 |
| F1_MAG_3 | 99.16 | 1.14 | 2.4 |
| F1_MAG_4 | 97.23 | 0.36 | 6.7 |
| F1_MAG__5 | 100 | 0 | 29.5 |
| F1_MAG_6 | 99.75 | 0 | 7.7 |
| G10_MAG_1 | 63.79 | 3.45 | 4 |
| G10_MAG_2 | 75.02 | 0.18 | 15.4 |
| G10_MAG_3 | 91.45 | 0 | 12 |
| G10_MAG_4 | 95.9 | 2.17 | 11.5 |
| G10_MAG_5 | 63.69 | 0.44 | 3.8 |
| H9_MAG_1 | 98.45 | 0.05 | 14.3 |
| H9_MAG_2 | 99.09 | 0.01 | 25 |
| H9_MAG_3 | 86.25 | 0.95 | 3.4 |

Table S6. Amino acid sequences of orthogroups predicted by OrthoFinder having the potential of TOrC biotrasformation.

| Orthogroup | Sequences |
|---|---|
| OG0016145 | >C7_bin.1_contig_14_2796<br>MCGIFGIAADSKKISQDQVAAALSSLFILSETRGKESAGITLVLPDEIVVQKSAVRPKQF<br>IRSHSYRQLIDQAMHRSRSAGTSYIAFGHSRLVTNGSQTVHANNQPVISGGIVGVHNGII<br>VNDDELSRTVPGIERRAAVDTEVFLQLVRLSMRAGRNVDDALRSAFNKVVGTVSTALVFE<br>DLDVLALGTNNGSLYVASAPDAGIFAFASESYILDSFVSERSLPWPTAENLIRPIQAKEC<br>LLVNLADRSLTSFHVDGAEQIGDLPHHRRNVRDDVPADENEMHGFDSAPAINHLNLSRYD<br>IDAEPIRRLRRCTRCILPETLPFIKFDANGVCNFCADYTPMKYRGRAALEIKAQQLRERR<br>RGTRADGLFTLSGGRDSSYGLHYAVRELGLKPVAYTYDWGMITDLARRNQARMCGALKVE<br>HILISADISKKRENIRRNVSAWLKKPSLGMIPLFMAGDKQYFYFANDLGKKLGLNTTILA<br>SNPLEKTHFKAGFCGVAPTQSHRPSKTAQIKMASFYAAHFLRNPAYLNPSLIDTVSAFGS<br>YYMISHNYLRLFDYVPWEEETINSTLLQEYDWETAPDTTSTWRIGDGTAAFYNYIYYRVA<br>GFSENDTFRSNQVREGAMTRTRALEMAETENRPRFESIAWYCSVIGLDPVDVLEKIAQIP<br>LLYPPR*<br>>C7_bin.2_contig_8_1885<br>MRDDDHYVVFRADDALTKLRRRVGLEGRQLFFGHSRLITNGLADNQPVVRGDVCVIHNGI<br>VVNHDQLWATIDKSPELEIDTEVIAAIAEWHLEQARPLEELGESVFGLCQGIVACAIAVP<br>RLGKLVLLSNNGSLYLGSKPNGIVFASESYPLTSMGCENVRQIRDAVVIDIPIDTDPVAV<br>KDWRGRKTNLVPALVLSSVEENLLEHPQPDLQRCTRCILPHTMPFIRFDADGVCNYCQNY<br>KPRNIPRPREELFELVEPYRRPGETDCIVPFSGGRDSCYALHLIVNELGMRPVTYTYDWG<br>MVTDLGRRNISRMSAELGVENIIVADDIWKKRDNIAKNLRAWLKSPHLGMVSILTAGDKH<br>FFRHVETMKRQTGIGLNLWGINPLEVTHFKAGFLGVPPDFEEERVYTHGAMKQLRYQYLR<br>LRAMAQSPGYFNSSLWDTLSGEYYRSFTEKSDYFHIFDYWRWDESTIDETLEAYDWERAP<br>DTQATWRIGDGTAAFYNYIYYTVAGFSEHDTFRSNQVREGDLTREEAIDLVKVENAPRYP<br>NIKWYLDAIGLEFEPVIKTVNAIPRLY*<br>>C9_bin.2_contig_204_378<br>MCGIFGIAADSKKISQDQVAAALSSLFILSETRGKESAGITLVLPDEIVVQKSAVRPKQF<br>IRSHSYRQLIDQAMHRSRSAGTSYIAFGHSRLVTNGSQTVHANNQPVISGGIVGVHNGII<br>VNDDELSRTVPGIERRAAVDTEVFLQLVRLSMRAGRNVDDALRSAFNKVVGTVSTALVFE<br>DLDVLALGTNNGSLYVASAPDAGIFAFASESYILDSFVSERSLPWPTAENLIRPIQAKEC<br>LLVNLADRSLTSFHVDGAEQIGDLPHHRRNVRDDVPADENEMHGFDSAPAINHLNLSRYD<br>IDAEPIRRLRRCTRCILPETLPFIKFDANGVCNFCADYTPMKYRGRAALEIKAQQLRERR |

| | |
|---|---|
| | RGTRADGLFTLSGGRDSSYGLHYAVRELGLKPVAYTYDWGMITDLARRNQARMCGALKVE<br>HILISADISKKRENIRRNVSAWLKKPSLGMIPLFMAGDKQYFYFANDLGKKLGLNTTILA<br>SNPLEKTHFKAGFCGVAPTQSHRPSKTAQIKMASFYAAHFLRNPAYLNPSLIDTVSAFGS<br>YYMISHNYLRLFDYVPWEEETINSTLLQEYDWETAPDTTSTWRIGDGTAAFYNYIYYRVA<br>GFSENDTFRSNQVREGAMTRTRALEMAETENRPRFESIAWYCSVIGLDPVDVLEKIAQIP<br>LLYPPR*<br>>D8_bin.5_contig_117_6<br>MIESLLAVAVLRDAPSSPQNRRNPVCGIFGYVGPDALDSSSTNTLVKHAQQRGRDSSGMV<br>MLGADGYHAYRADYRIGWLLKRIPKPTNLFFGHSRLVTNGTGDNQPVLREQVLVLHNGIV<br>VNEEQIWTSFGKKPGSPSTPRSSPRSSPPISPPAARSRPPPSGCSSSPSASSPARPSSRR<br>SASSCCSRTTAASTSPTSPAAPCSRRSATRSSRSGRRTSARSSTTSRSSTCRWRRRRSRS<br>PSGRTAPSTSSPASA*<br>>D8_bin.5_contig_117_7<br>MPFSGGRDSSYGLHLIIHELKLRPVTYTYDWGMVTDLGRRNVSRMSSMLGVENIIVAADI<br>TTKRDYIRRNLEAWIKRPHLGMLSVLTAGDKHFFRHIETLKRQTGVSLNLWGINPLEVTH<br>FKSGFLGVPPDFAEERVYSHGALKQLRYQSLRFRAMLQSPGYFNRSIPDTLLGEYYRSFT<br>TKSDYFHIFDYWRWDEKLIEETLDAEYDWEHAPDTQTTWRIGDGTAAFYNYAYYTIAGFT<br>EHDTFRSNQIREGDITREEALRLVHEENTPRYPNLKWYLDVVGVDFTKAITAINRAPRLY<br>EGAPHPSFAAPRD* |
| OG0023482 | >C7_bin.1_contig_14_3102<br>MSTTE-----KAVSTSDALVAATKDESVHHIVVQGNLTGAPTINLLPGQSLRGDGDAATI<br>SFAKGSDGLRLSSDNRIHNIRLNTAVEKRAIFNDTSVDSFGRIELRGVITTGRVQILARD<br>KVRGGHVDVIGLDIVAADARAEMDRPQGYGVYVLQGAFTLWNMQQDTNVTISADLVGLSA<br>GRDGAPVRGSGIFVSGGGDKAGKLAVRRLETDAVYSDGGIASGTPDQITGGVFTVYGAHV<br>DVVRNRGPVTTYGVNDMVLDNWGVVDRWTAEAKITSHGPSGICFVNFGIVHELKVNAPIE<br>TFGQGARGFNVYTGTVNLAEFDRVTTHADGSVGIQISQPIGRLVVHRGIETFGGTGPSLV<br>KGKVITLSAIGLSVKPGGSAREIEISGGIKTNGAGVSPIEQHGAIEKLSVSGGFVAAGGG<br>FDKI*<br>>C9_bin.2_contig_204_73<br>MSTTE-----KAVSTSDALVAATKDESVHHIVVQGNLTGAPTINLLPGQSLRGDGDAATI<br>SFAKGSDGLRLSSDNRIHNIRLNTAVEKRAIFNDTSVDSFGRIELRGVITTGRVQILARD<br>KVRGGHVDVIGLDIVAADARAEMDRPQGYGVYVLQGAFTLWNMQQDTNVTISADLVGLSA<br>GRDGAPVRGSGIFVSGGGDKAGKLAVRRLETDAVYSDGGIASGTPDQITGGVFTVYGAHV<br>DVVRNRGPVTTYGVNDMVLDNWGVVDRWTAEAKITSHGPSGICFVNFGIVHELKVNAPIE |

| | |
|---|---|
| | TFGQGARGFNVYTGTVNLAEFDRVTTHADGSVGIQISQPIGRLVVHRGIETFGGTGPSLV<br>KGKVITLSAIGLSVKPGGSAREIEISGGIKTNGAGVSPIEQHGAIEKLSVSGGFVAAGGG<br>FDKI*<br>>D8_bin.3_contig_1_313<br>MSSSETAANMKSVTTVDELVAATKDKSAHHIVVRGGLTNAPSIRLAPGQSLRGDGDTAAI<br>TFAAGIDGLQLSSDNRVHNLRLNATVNKRAIFNDTSVESLGRIELRSVTTTGCVQILARD<br>KVRGGHVDVNGLDIIAADARGEQERPHGYGVYVINGAFTLWNMQQDTGVVVSADLVGLSA<br>GRDGAPVRGSGIFVSGGGDKAGRLNVRRLETDAVYSDGGIAPGTADQITGGVFTVYGAYV<br>DVVRNRGPVVTYGVNDMVLDNWGVVDRWTAEAKITSHGPSGIGFVNFGIVHELKVNAPIE<br>TFGQGARGFNVYTGTVNLAEFDRVITHADGAVGVQISQPIGKLVVHRGIETFGGTGPSLV<br>KGVVITLSAIALSIKPGGSAREIEISGGVKTNGAGVAPIEQHGAINSLRVTGGFVAASGG<br>FDKI* |
| OG0019876 | >A10_bin.3_contig_1_5674<br>MKNRVFRFLTTATLAVSVALAPVAFARGGGGGGHGGGGGHGGGFGGGMHGGGMAFAAMGG<br>GGHFTGGHFGGARFAGAGAFGPRFAGAGFHGARFFHHGGFFHHRFHRFAFFGAPYLYAGY<br>DDGCWRRAWTSYGLQWVNVCGDYW*<br>>A2_bin.4_contig_123_1961<br>MNRRILTGLLAAVLATTVALASPAFARGGGYGGGGHGGGFHGGGGWHGGGGMRAMGGGMR<br>FSGIGGGPRFAGARIAGPRFAHAGIHRGFHHRFHHRRFAFIGAPYLYAGYNYGCWRRVWT<br>GYGPRWVNFCDYGYGLY*<br>>G10_bin.1_contig_2_1812<br>MAGRGVVAPGGHRFVGAPFAGRQAFTHQFHRGFAFRHHHRFNRFVFVGTPFAYASYAAYD<br>GCWREVWTGYRWRWANTCNGYGYGYGY* |
| OG0019848 | >A10_bin.3_contig_1_2298<br>MSRGKRRKVSIAVRRVALAKQKLGRDQLKRFKDWDDGRPETELNFKFYRQATNKIVGITD<br>ETAAFLEGLF*<br>>A2_bin.2_contig_237_4099<br>MIFAIKAEVNDPRAKTLEFSPQKTMYGGKLIAPGDTIFVFASENEGGPGLIASGIVTSTQ<br>TVARKRGVARQTPRVSVTIRRTALAQRRLGRSELKLFSSWNDGRPETELNFKFYRQATNK<br>IVGLSEAAAAFLGEFF*<br>>G10_bin.4_contig_320_406<br>MAFAIKAEIADPRAEHFSFAAQKTMYGGKNIAVGDTIFLFASENDGGRGLVARGIVLSAQ<br>EVGRQPDIERQTPRVSITVRRTAFARRPLGRNELRHLTDWQDGRPGTELNFKFYRQATNK<br>IIGISDDASMFLKSFF* |

| | |
|---|---|
| OG0031888 | >B4_bin.1_contig_3_202<br>MARSIRLLYRSQHGTIRKNFNWDPINLNSTVIITAAEFTPAFGGLGGGPKTLGRPNLGLA<br>NVYVTNVGPHGRAGVEAGGVEFLLHVDWNSPLDIVVTITVLDDIEQFVQA*<br>>F1_bin.2_contig_13_380<br>MSNSVRWVIRGIRGRVPANFNWGIISARSIVHVSAAEVRFGTTQVNPQPPLQNFFYNLGA<br>ADIWVSNISPHRNEFSGQPGGVSFIVHVGWNSPLDVAVTITVEDALPVEIQGY* |

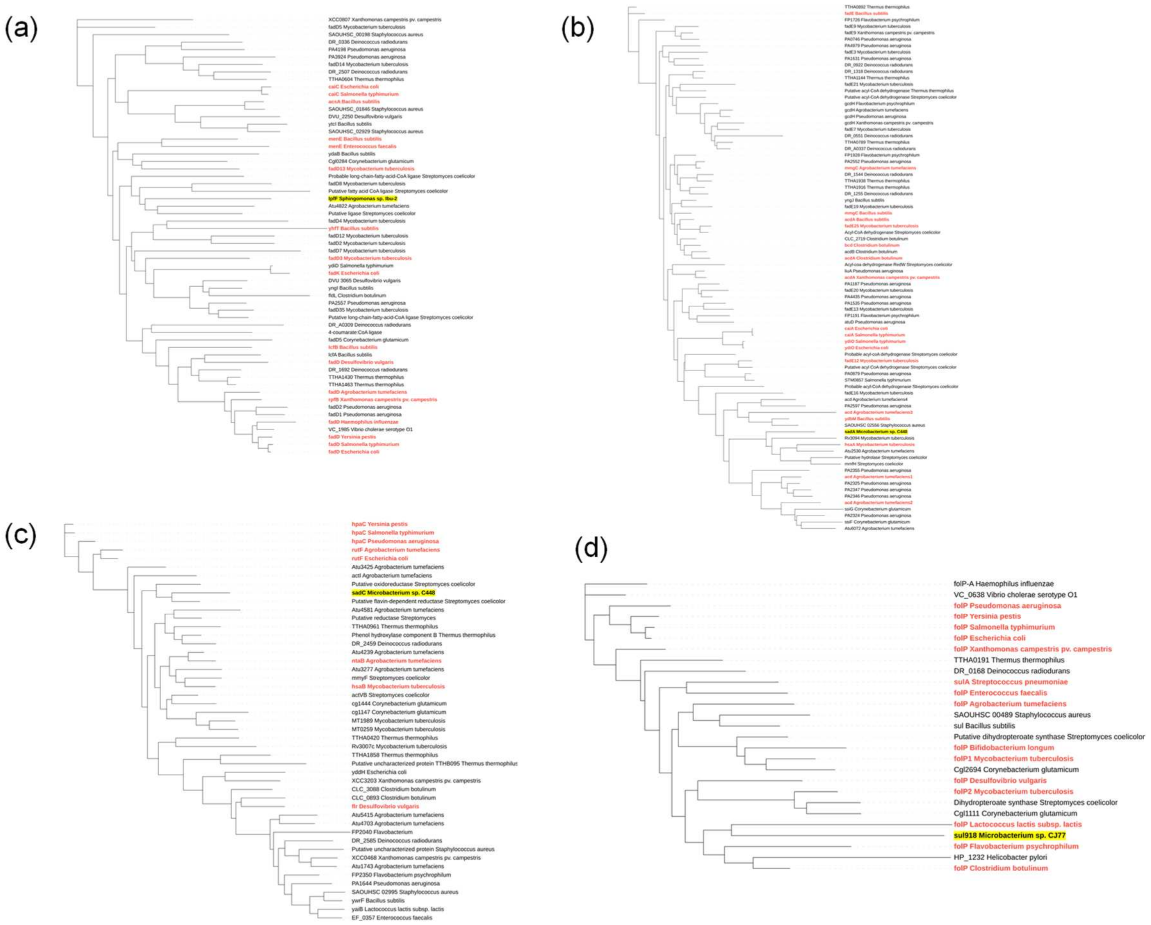


Figure S1. Phylogenetic trees of (a) *ipfF,* (b) *sadA,* (c) *sadC,* and (d) *sul918* genes and their homologs. Homologous proteins were identified based on the amino acid sequences of these biotransformation genes using OrthoFind. Phylogenetic trees were constructed using maximum likelihood method by IQ-TREE and modified by iTOL. Red color represents the gene existence in the genomes of model communities.

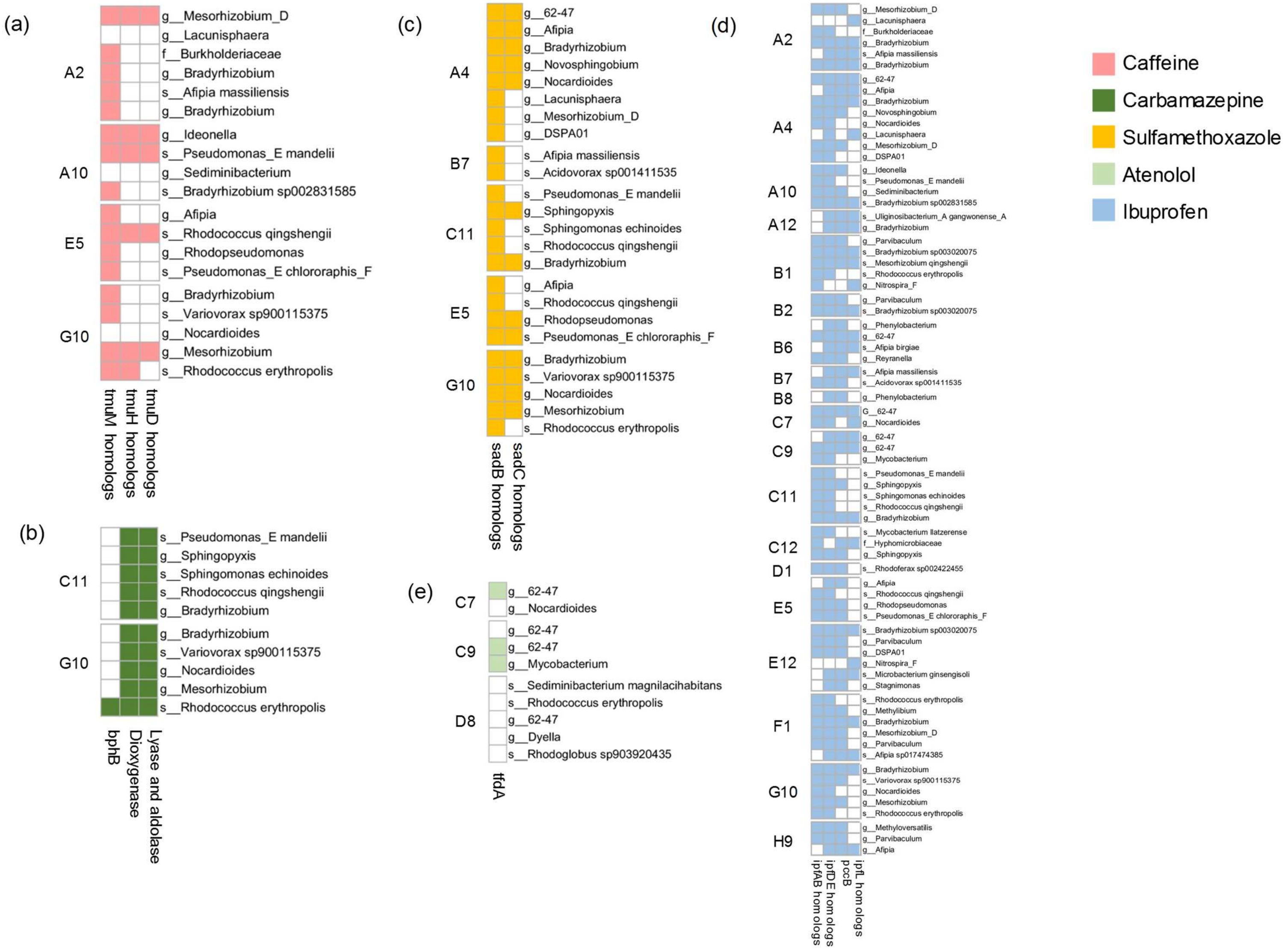


Figure S2. Presence of biotransformation genes and enzymes and their homologs of the subsequent reactions after the first metabolic step in degrading model communities. Colors represent presence, blank represents absence.
